# Forward Engineering Organ Development and Cancer Therapeutics with Optogenetics

**DOI:** 10.1101/2025.05.16.653804

**Authors:** Mayesha Sahir Mim, Stephen Cini, Caitlin Frank, Zian Wang, Alexander Dowling, Jeremiah J. Zartman

**Affiliations:** Chemical and Biomolecular Engineering, University of Notre Dame, Notre Dame, IN 46556, USA; Bioengineering Graduate Program, University of Notre Dame, Notre Dame, IN 46556, USA

**Keywords:** CsChrimson, Organogenesis, Apoptosis, Compensatory Cell Proliferation, Ras^V12^, *Drosophila*

## Abstract

Robust growth control is an essential requirement for the survival of living organisms, while its dysregulation results in diseases such as cancer. However, a significant knowledge gap exists in understanding how precise organ growth control is achieved. The growing arsenal of optogenetic toolkits allows precise, noninvasive control of cellular signaling in vivo, enabling research into how bioelectrical and chemical cues regulate organ growth. Here, we used the red-light-activated channelrhodopsin, CsChrimson, to stimulate intracellular calcium signaling dynamics in the wing epithelium of *Drosophila melanogaster*, an established model system for investigating organ size control. By varying light intensity and activation dynamics systematically, we identified a biphasic regulation of final organ size. Illumination of CsChrimson depolarizes cells and stimulates spikes of cytosolic calcium concentrations, a phenomenon explained by a computational model that incorporates the inclusion of both gap junction closure and voltage-gated calcium channel activation. This calcium regulation tunes downstream effectors involved in growth regulation and apoptosis. In particular, we found that prolonged bright red light exposure (100 lux/12 hours) increased cell death in wing imaginal discs and caused severe morphological abnormalities in adult wings, with phenotypic severity dependent on stimulation parameters defined by illumination intensity and period of activation. Strikingly, an optimum level of dim, pulsatile light (5 lux, 1 minute on/off pulse train) resulted in overgrown organs and significantly upregulated cell proliferation. We also co-expressed an oncogene, *Ras^V12^*, with CsChrimson and showed that experimental optical simulation parameters can be exploited to control the morphology of tumorous tissues and initiate targeted remission of tumorous growth. Our findings and approach provide a powerful framework to dissect the role of dynamic physiological signaling events in organogenesis and offer translational insights into new therapeutic strategies with applications in cancer and regenerative medicine.

## Introduction

During organ development, the spatial and temporal distribution of morphogens provides critical positional cues that establish cell identities across undifferentiated tissues^1,2^. These signals, along with other chemical gradients, mechanical forces, and electrical fields, orchestrate morphogenesis, an inherently multiscale and multilevel phenomenon involving the three-dimensional shaping of tissues over time^3,4^. Precise control of this morphological control is essential for understanding developmental disorders, tissue repair, and regenerative medicine. Integrating computational and experimental approaches across diverse systems is key to uncovering how genetic and biophysical mechanisms govern organ growth, as it provides a systems-level quantitative and predictive understanding of morphogenesis. This motivates the use of mathematical models grounded in a holistic biophysical approach that captures both molecular signaling and emergent supracellular dynamics^5,6^. The complex, multimodal nature of organogenesis, spanning chemical, mechanical, and biophysical factors, remains a significant challenge for bioengineers to unravel, yet solving this mystery holds promise for advancing tissue engineering, organ regeneration^7^, and targeted therapies for human diseases such as cancer.

Optogenetics offers a powerful means of controlling cellular processes with high spatial and temporal resolution^8–10^. Optogenetic technology has advanced into therapeutic applications, with clinical trials reaching Phase 3 for treating retinitis pigmentosa, and preclinical studies demonstrating its potential in modulating neural circuits in models of schizophrenia and epilepsy-induced seizures. By expressing light-gated ion channels, such as channelrhodopsins^11^, in transgenic animals, specific biochemical activities can be activated with light to modulate subcellular processes and tissue development. Upon activation, these channels facilitate the influx of cations, including calcium (Ca²⁺), sodium, potassium, and hydrogen ions, triggering intracellular cascades critical for morphogenesis. CsChrimson^12^, a red-shifted, plasma membrane-localized channelrhodopsin, enables deep-tissue light penetration and dual-color experiments, extending optogenetic utility into epithelial biology^13^. While well-established in neuroscience^7–11^, the application of optogenetic channels, like CsChrimson in epithelial tissues to particularly in tissue and organ developmental contexts remains understudied.

Ion signaling is a key regulator of cell growth. Divalent ions such as Ca²⁺ and magnesium (Mg²⁺), along with calmodulin, regulate both normal and neoplastic human cell proliferation^14^. In fungi and mammalian systems, the growth response to sodium (Na⁺), potassium (K⁺), and other cations further highlights the broader physiological role of ion signaling^15,16^. Ca²⁺, in particular, serves as a versatile second messenger, orchestrating cell proliferation, apoptosis, and migration, and is regulated by Ca²⁺-selective channels that mediate influx from the extracellular space into the cytosol. Different categories of Ca²⁺ signaling, spikes, transients, waves, and fluttering, translate upstream inputs into coordinated multicellular responses. For instance, “initiator” cells with elevated phospholipase C (PLC) activity, as opposed to “standby cells” with baseline PLC activity, trigger intercellular Ca²⁺ waves via Gαq signaling, while insulin signaling can induce localized Ca²⁺ spikes^17–19^. These dynamics also influence tissue mechanics, as the spatial context of signaling events modulates mechanical responses during development^20^. Despite these insights, the current understanding of biological mechanisms precludes direct control of Ca^2+^-mediated downstream processes, impeding the understanding of its relation to organ growth. Previous studies have explored the Hedgehog pathway and the role of cations in stimulating the release of morphogens, such as bone morphogenetic proteins in regulating final organ size^21–24^. However, Ca²⁺ signaling is increasingly recognized as an essential regulator of organogenesis^25–27^. Disruptions in Ca^2+^ signaling can lead to developmental abnormalities^28^, underscoring the need for precise control of Ca²⁺ signaling dynamics in vivo, yet its downstream influence on tissue growth remains poorly understood.

The wing imaginal disc, a model epithelial tissue of *Drosophila melanogaster* is a well-established system to study morphogenesis and gene function in epithelial tissues as the wing and its primordial wing imaginal disc are acutely sensitive to changes in developmental pathways^29,30^. Its sensitivity to genetic perturbations due to reduced functional redundancy in paralogous genes found in vertebrate systems, along with powerful tools like the Gal4/UAS system^31^, enables spatially precise manipulation of gene expression (**Figure S1**). This makes it a valuable system to explore how subcellular signals scale to tissue-level outcomes such as organ size and shape. Furthermore, the conservation of core Ca²⁺ signaling pathways between *Drosophila* and vertebrates facilitates broader relevance. *Drosophila* and the wing disc model has also enabled the *in vivo* study of Ras^V12^, a constitutively active mutation in the rat sarcoma virus (RAS) protein family that drives tumorigenesis, highlighting conserved mechanisms such as Piezo-mediated invasion and NF-κB signaling in tissue homeostasis and cancer^32–34^. Although optogenetics is widely used in *Drosophila* neuroscience^35,36^, its application in epithelial morphogenesis, particularly to dissect Ca²⁺-mediated tissue growth and neoplastic transformation, remains underexplored.

In this study, we introduce an optogenetic approach to directly manipulate cytosolic Ca²⁺ dynamics and uncover their role in growth-related signaling pathways that govern proliferation and apoptosis. Using the red-light-activated channelrhodopsin CsChrimson, we generated transgenic *Drosophila* to precisely control intracellular Ca²⁺ levels in the developing wing epithelium. Activation of CsChrimson-expressing cells stimulated intracellular Ca²⁺ dynamics. To better understand the underlying mechanisms, we developed a computational model that recapitulates these experimental findings. The model demonstrates that resulting Ca²⁺ dynamics can be explained by simulating gap junction closure and subsequent activation of voltage-gated calcium channels as key drivers of the observed calcium fluctuations. By integrating experimental data with computational modeling, we identify new mechanistic insights into how optogenetically induced electrical changes propagate through tissue networks and modulate Ca²⁺ signaling at the multicellular level. Our findings show that prolonged optogenetic activation induces severe organ-level phenotypes, marked by increased apoptosis, reduced mitosis, and downregulation of integrin expression. In contrast, stimulation within a defined, low-intensity activation threshold promotes optimal organ growth by reducing apoptosis and enhancing compensatory proliferation in the growing tissue. Based on these insights in robustly developing tissues, we also demonstrate that this light-controlled system can be harnessed to induce remission in *Ras*^V12^-driven tumorigenesis, highlighting its potential as a tool for modulating aberrant growth. Together, our results establish a direct role for Ca²⁺ and cationic signaling in organ development and offer a tunable platform for probing growth regulation *in vivo*.

## Results

### CsChrimson Triggers Spikes in Calcium Concentrations in Non-Excitable Cells

Genetic expression of the synthetic optogenetic tool, CsChrimson, stimulates Ca²⁺ and other cation transportation in *Drosophila* neurons through a mechanism of voltage-gated Ca^2+^ channels^37–40^, where the channel stays inactive in darkness and optically activates to induce ion flux with red light illumination (**Figure 1A**). We hypothesize that this holds true in developing epithelial tissues like the wing imaginal disc, and by systematically controlling light duty cycles, growth-related processes can be directly modulated in the cells expressing CsChrimson and other ion transporters, developing tissues and eventually, terminal organs (**Figure 1B**). Building on previous findings that Ca²⁺ signaling dynamics can directly tune tissue and organ development through hormonally driven and gap junction–mediated activity patterns in the *Drosophila* wing disc^17^, we investigate how an artificial light-gated channel can be used to externally modulate Ca²⁺ dynamics and influence developmental signaling.

**Figure 1:**
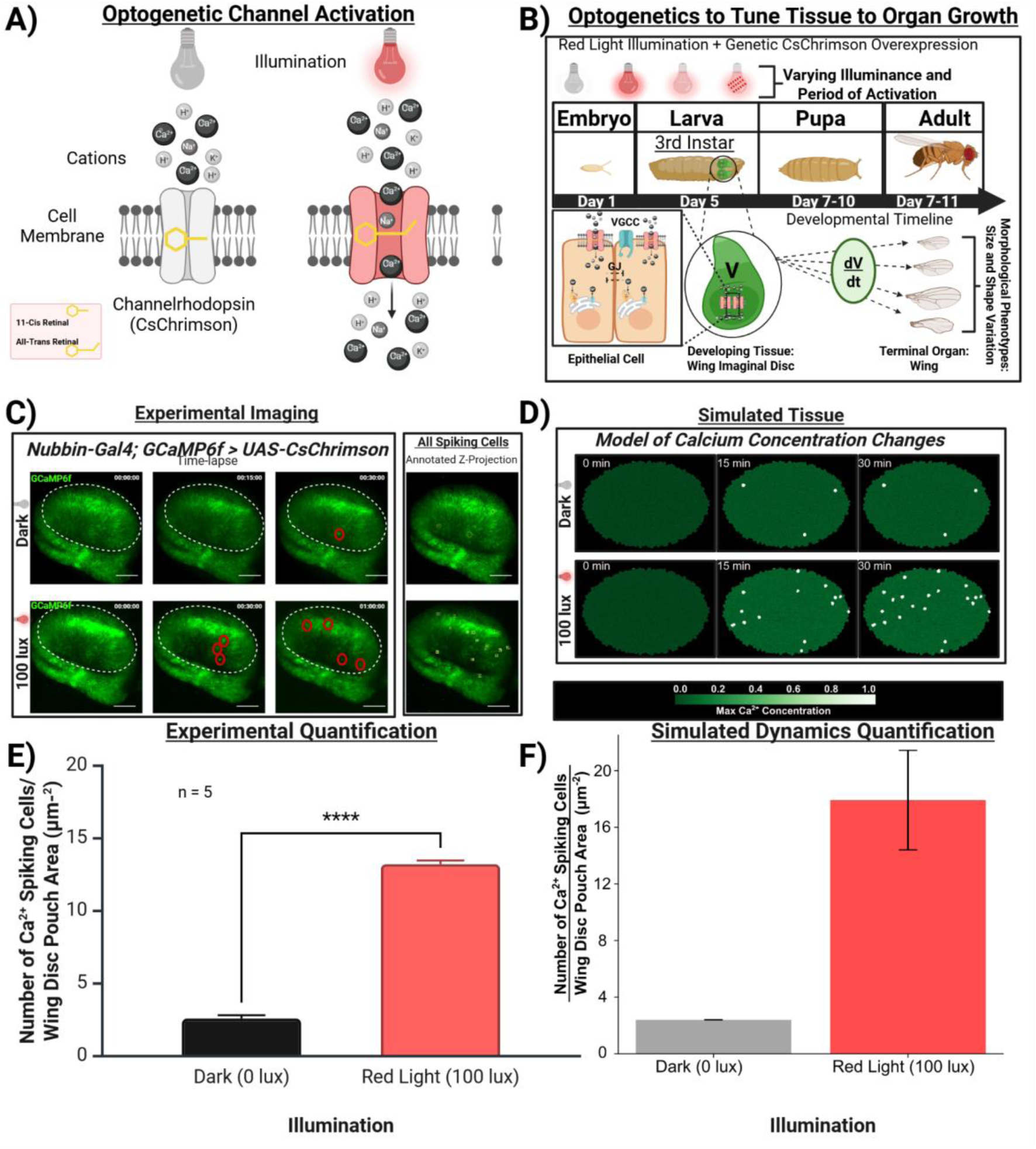
Activation of CsChrimson Stimulates Ca^2+^ Spikes in the Wing Imaginal Disc. **A)** Schematic of CsChrimson’s inactive state in darkness and activation under red light. **B)** Schematic illustration of hypothesis: systematic light-controlled activation of CsChrimson can modulate growth-related processes in developing wing discs and the morphology of terminally differentiated adult wings. Within the epithelial cells, GJ = Gap Junction (inactivated), and VGCC = Voltage Gated Calcium Channel. **C)** Representative experimental images extracted from 30 min/1 min imaging at 0, 15 and 30 minutes, and a max projection over time under dark (top panels) and 100 lux red light illumination (bottom) for a wing imaginal disc overexpressing CsChrimson in the wing disc pouch driven by a Nubbin-Gal4 driver tagged with GCaMP6f. Red and yellow circles indicate the occurrence of Ca^2+^ spikes. Scale bar = 50 µm. **D)** Simulated wing disc showing elevated Ca^2+^ activity. Top row: Dark simulation at timepoints = 0, 15, 30 minutes; Bottom row, red light activation at timepoints = 0, 15, 30 minutes. Scale on the right side shows Ca^2+^ concentration. **E)** Bar plots quantifying the number of experimentally observed Ca^2+^ spiking cells per area of the wing disc pouch over the period of 30 minutes. Unpaired t-test was performed. Four stars (****) signify p-value less than 0.0001 **F)** Similar quantification in computationally modeled simulated tissue.

To test whether CsChrimson can depolarize non-excitable *Drosophila* tissue, we used the *Nubbin-Gal4* driver to express *UAS-CsChrimson* specifically in the wing disc pouch and activated CsChrimson using 600 nm red light using a customized optogenetic activation protocol to determine whether discernible cytosolic Ca²⁺ dynamics could be observed (**Figure S1, S2, S3**). The wing imaginal discs were dissected from third instar larvae reared on food supplemented with 1 mM all-trans Retinal to ensure functional activation of the optogenetic channel^41^. A genetically encoded Ca^2+^ indicator, GCaMP6f^42^, combined with the *Nubbin* driver, enabled real-time visualization of cytosolic Ca²⁺ dynamics.

As a control, the wing disc was imaged in darkness every minute for 30 minutes. Under these conditions, minimal Ca²⁺ activity was detected (**Figure 1C**, top panels). However, upon optical activation of the same sample using 100 lux red light, Ca²⁺ spikes were stimulated nearly instantaneously and were sustained over the same period of imaging (**Figure 1C**, bottom panels), as shown in the annotated projection over time. Wing disc pouch cells exhibited increased Ca²⁺ spiking activity (**Video S1)**. Quantification of the total number of spiking cells over each wing disc pouch area confirmed that cytosolic Ca²⁺ spikes increased dramatically upon optical activation, while overexpression of CsChrimson alone in the absence of light had no significant effect on Ca²⁺ dynamics **(Figure 1E)**. These results confirm that CsChrimson can be functionally activated in non-excitable epithelial wing disc cells and reliably induces sustained Ca²⁺ spiking upon red-light activation.

To mechanistically interpret how red-light stimulation of CsChrimson, which depolarizes cells^12^, drives tissue-wide Ca^2+^ responses, we adapted a previously established computational model^17^ of calcium signaling to simulate and test whether addition of a disturbance term can explain the calcium signaling dynamics observed during experimental optogenetic activation (**Figure 1 D, F,** <u>Methods</u> and Supplementary Information). This model captures tissue-scale Ca^2+^ behavior in epithelial cells as a function of an increased optogenetic stimulus. To integrate optogenetic control into our calcium dynamics model, we introduced a light-dependent control term, u(t), directly into the differential equation describing cytosolic Ca^2+^ dynamics, with equations, assumptions, and parameter table detailed in the Materials and Methods section. In this model, optogenetic activation of CsChrimson does not induce Ca^2+^ dynamics directly. Instead, the ’opto.term’ u(t) estimates the direct addition of Ca^2+^ concentration resulting from CsChrimson-mediated membrane depolarization. Interestingly, initial simulations that only incorporated a Ca^2+^ flux into cells led to intercellular Ca^2+^ transients (**Figure S4 A)**. This prompted a second scenario that also incorporates depolarization-dependent gap junction closure in the model. This approach reflects experimental evidence showing that CsChrimson activation induces immediate increases in intracellular calcium through voltage-gated calcium channel activation and depolarization-dependent gap junction closure^43–46^. Thus, simulations and literature support in other studies collectively suggest that the light-dependent Ca^2+^ dynamic responses observed in our experimental system likely is through voltage-gated calcium signaling and gap-junction regulation. Future studies will require identifying which voltage-gated channel is involved. Thus, computational modeling coupled to experimental observations provide insights into how bioelectrical and intercellular signaling are integrated during optogenetic stimulation. (**Figure 1 C-F)**.

Additionally, our simulation results show that at a 100 lux red light illumination, corresponding to an optogenetic scaling factor of ɑ = 0.14, Ca^2+^spiking increased in the original initiator cell, which already exhibited Ca²⁺ dynamics at 0 lux (**Figure S4 B)**. Ca^2+^ spiking was newly induced in additional bystander cells. Simulations across a range of ɑ values from 0.05 to 1 demonstrated how increasing the input light stimulus, u(t), influences the fraction of initiator cells: the fraction begins to rise at ɑ = 0.139 and saturates, reaching a fraction of 1, at ɑ = 0.34 (**Figure S4 C-D)**.

Together, our *in vivo* and *in silico* experimental findings highlight the role of optogenetically induced Ca²⁺ signaling, while our computational model provides mechanistic insight into the spatiotemporal dynamics of Ca²⁺ activity during development.

### High-Illuminance CsChrimson Activation Disrupts Wing Development

Given the critical and multi-faceted roles of Ca²⁺ signaling in modulating *Drosophila* wing organogenesis^17,18^, we next sought to investigate how the CsChrimson-induced increase in cytosolic Ca²⁺ within the wing disc impacts the development and morphology of the terminal adult wing. To test whether CsChrimson-mediated elevated cytosolic Ca²⁺ levels at the tissue level could directly impact the mature organ, we again expressed *CsChrimson* specifically in the pouch region of the wing imaginal disc using the *Nubbin-Gal4* driver. This targeted expression was restricted to the developing wing, allowing us to observe adult wing phenotypes without perturbing other tissues or organs that are critical for overall animal viability. Optical activation with 100 lux red light, delivered continuously over 12 hours from the first through third larval stages, resulted in a range of severe wing phenotypes in F1 adult progeny, including complete wing loss, shrivelled wings, blisters, and failure of wing expansion (**Figure 2A**).

**Figure 2:**
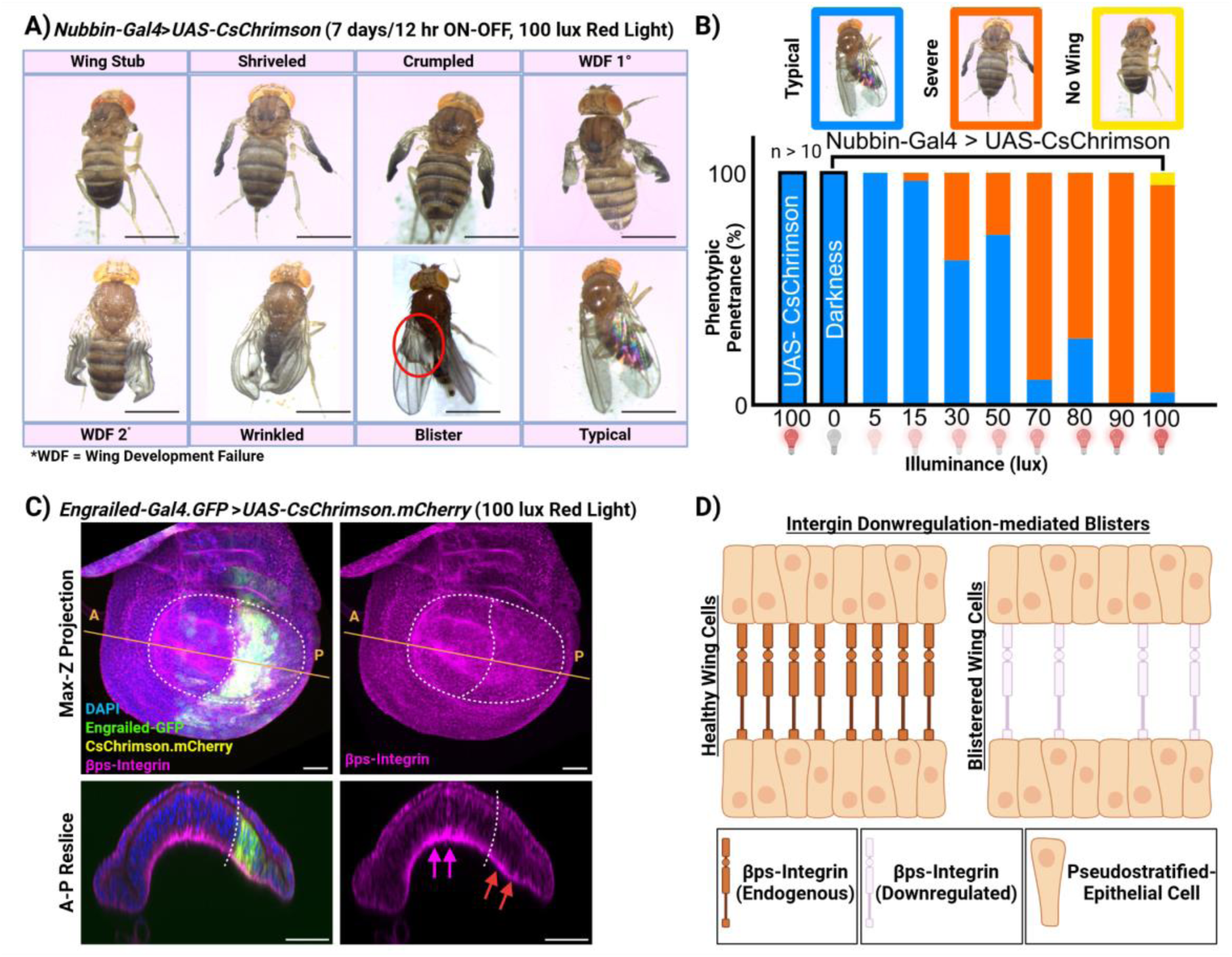
Enhanced tissue-level optogenetic activation causes severe organ phenotypes. **A)** Multiple severe phenotypes diverging from regular wing morphology emerge when CsChrimson is activated in the wing imaginal disc, including missing wings (wing stub), asymmetric left and right wings, shriveled and crumpled wings, failure of robust and complete wing development at multiple stages wing development failure (WDF), and blistered wings. Red circle highlights the blister on the wing. Scale bar = 1 mm. **B)** Bar chart quantifying wing phenotypes emerging at gradually increasing illuminance levels of red light from 0 lux to 100 lux. Two leftmost bars signify controls (*UAS-CsChrimson* = Parental control without genetic perturbation, and Darkness = Genetic overexpression of *CsChrimson* without optical activation). Blue = morphologically typical *Drosophila* wings, Orange = wings with severely damaged phenotype, Yellow = flies with only wing stub and no fully formed wing. Sample sizes corresponding to each bar are as follows: n_UAS-CsChrimson/100lux_ = 72, n_Perturbation/0lux_ = 50, n_Perturbation/5lux_ = 23, n_Perturbation/15lux_ = 83, n_Perturbation/30lux_ = 90, n_Perturbation/50lux_ = 109, n_Perturbation/70lux_ = 28, n_Perturbation/80lux_ = 12, n_Perturbation/90lux_ = 11, n_Perturbation/100lux_ = 97. **C)** Maximum Z projection and corresponding optical reslices across the anterior (A)-posterior (P) axis along the orange line. White dotted boundaries show the shape of the wing disc pouch as the region of interest and the boundary of the *Engrailed* expression region. Fluorescent labels are stated on the bottom left. Scale bar = 50 µm. n = 8. **D)** Schematic relating the downregulation of integrin leading to blister-causing loss of adherence between dorsal and ventral epithelial cell layers in the Drosophila wing disc.

To assess how varying light intensities influence the organogenesis process, we activated CsChrimson using a range of red light intensities: 0 lux (dark), 5 lux, 15 lux, 30 lux, 50 lux, 70 lux, 80 lux, and 100 lux. The corresponding irradiance values are provided in Supplementary Information (**Table S1**). Two controls were used: (1) the genetically inactive parental line exposed to 100 lux red light and (2) flies overexpressing CsChrimson but reared entirely in the dark. Both controls were developed with morphologically normal adult wings (**Figure 2B, two leftmost bars).** In contrast, flies exposed to red light showed a clear dose-dependent relationship between illumination intensity and the severity of wing phenotypes. At low light intensity (5 lux), no morphological abnormalities were observed. However, increasing the light amplitude over the same seven-day developmental window during larval growth progressively led to more severe phenotypes. At 100 lux, approximately 90% of F1 progeny displayed severe wing defects, with some individuals emerging with only wing stubs. These results demonstrate that optogenetic activation of CsChrimson modulates morphogenetic outcomes in a light-intensity-dependent manner, underscoring the direct link between elevated Ca²⁺ and cation signaling and disrupted wing development.

The blister phenotype in the *Drosophila* wing is known to result from the loss of βPS integrin, which disrupts cell–cell adhesion and leads to detachment between the dorsal and ventral wing surfaces^47^. To determine whether this mechanism also underlies the phenotypes observed with 100 lux red-light activation, we performed antibody staining for myospheroid (βPS integrin) in wing discs overexpressing *CsChrimson*. Expression was driven specifically in the posterior compartment using *Engrailed-Gal4*, leaving the anterior compartment genetically unaltered and serving as an internal control **(Figure S1)**. We observed a marked downregulation of βPS integrin expression in the optogenetically activated posterior region of the wing disc, specially on the basal side as seen on the optical cross-section **(Figure 2C)**. This is consistent with the known mechanism of integrin-led loss of adhesion contributing to the observed wing blistering **(Figure 2D)**. Interestingly, in addition to blistering, other phenotypes such as wing asymmetry and wing development failures have also been observed upon genetic perturbation of another calcium-transporting mechanosensitive cation channel, *Piezo*^48^, and knockdown of the Sarco/Endoplasmic Reticulum Calcium ATPase, *SERCA*^18,49^ This suggests that alterations in cytosolic Ca²⁺ levels can directly influence morphogenesis through distinct ion channels operating via similar mechanisms.

These findings demonstrate that sustained optogenetic activation leading to an elevation of cytosolic Ca²⁺ in the developing tissue can disrupt key morphogenetic processes, including adhesion and tissue integrity, resulting in a spectrum of severe adult organ phenotypes. However, beyond overt malformations, we next asked whether more subtle changes in wing morphology, such as differences in overall wing area, might also emerge under moderate levels of optogenetic stimulation. To address this, we examined adult wings that developed without gross defects to assess whether CsChrimson-induced Ca²⁺ elevation produces quantifiable variation in wing area.

### Optimal Balance of Calcium Levels is a Catalyst for Organ Growth Control

To further investigate potential subtle effects of CsChrimson-induced Ca²⁺ elevation, we measured the area of morphologically normal wings, which resembled those of wild-type fruit flies. For these phenotypically typical wings, the entire wing blade, derived from the wing disc pouch region, was analyzed using the wing phenotype analysis tool, MAPPER, to detect any non-qualitative variations in size.^50^

Surprisingly, both very dark or low illumination (0, 3 lux) and higher (10, 50 lux) levels of illumination resulted in smaller wing blade areas compared to an intermediate level of activation (5 lux), which produced significantly larger wings (**Figure 3A, C, E**).

**Figure 3:**
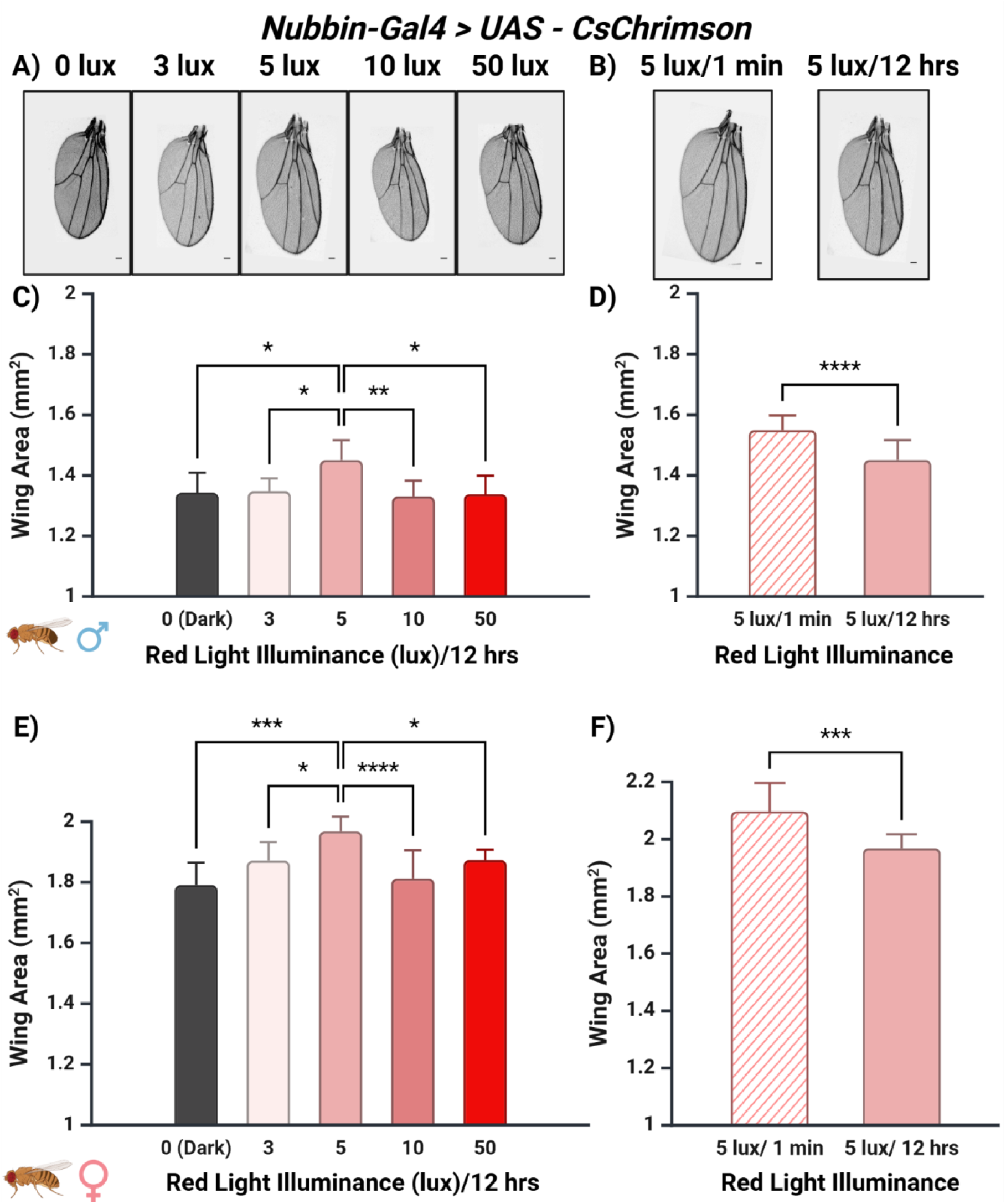
Optimal balance of Ca^2+^ levels is a catalyst for organ growth control. **A)** and **B)** Variation in wing size based on wing blade area for intensity and period of red light activation. Representative wings were taken from male samples. Scale bar = 100 µm. **C), E)** Bar plot showing the variation in wing area for both C) male and E) female fly wings for the optimum level of 5 lux, along with higher (10, 50 lux) and lower (0, 3 lux) illuminances. Colors on each bar reflect the illuminance level. Sample sizes from left to right, n_0lux,male_ = 6, n_3lux,male_ = 5, n_5lux,male_ = 11, n_10lux,male_=11, n_50lux,male_ = 7, and n_0lux,female_ = 6, n_3lux,female_ = 12, n_5lux,female_ = 9, n_10lux,female_ = 24, n_50lux,female_ = 9. **D), F)** Column plots of D) male and F) female wing areas at 5 lux/1 minute red light activation. n_5lux/1min,male_ = 30 and n_5lux/1min,female_ = 30 For C and E Kruskal-Wallis test with Dunn’s multiple comparisons test and for D and F Unpaired t test were performed. If p-value is less than 0.05, it is flagged with one star (*), less than 0.01 with two stars (**), less than 0.001 with three stars (***), and less than 0.0001 with four stars (****).

This phenotype size trend was consistent across all samples, showing no sexual dimorphism between smaller male wings and relatively larger female wings.

To further converge towards a larger organ using 5 lux illumination, we used a 1-minute on/off duty cycle, as this frequency activated the ERK1/2 (extracellular signal-regulated kinase) pathway^51^ which plays a crucial role in integrating external signals, such as epidermal growth factor, to regulate cellular processes, including proliferation. This stimulation strategy for growth signaling cascade activation proved successful, and we observed a further increase in wing area in both male and female fly wings as opposed to 5 lux illumination over 12 hours (**Figure 3B, D, F**).

Together, these findings demonstrate that fine-tuned optogenetic activation, specially at low light levels, can enhance organ growth by modulating intracellular signaling networks. Based on these insights at the organ level, we next focused on analyzing subcellular processes, specifically cell proliferation and programmed cell death, regulated by CsChrimson activation within the wing imaginal disc.

### Optogenetically-Tunable Cell Fate Shifts through Apoptosis and Compensatory Proliferation

Using antibody staining, we investigated how adult wing phenotypes correlate with growth-related mechanisms, by examining subcellular processes downstream of Ca^2+^ signaling including cell proliferation and programmed cell death. Immunohistochemistry was performed with antibodies Phospho-Histone H3 (PH3) to label mitotic cells and Death Caspase 1 (DCP1) to detect apoptotic cells in the wing imaginal disc^52,53^. Consistent with our previous optical perturbation and findings, we used a range of illumination intensities and activation duty cycles, including complete darkness, continuous 100 lux for 12 hours, 100 lux pulsed every 1 minute, continuous 5 lux for 12 hours, and 5 lux pulsed every 1 minute, to identify conditions most conducive to growth (**Figure 4A, S4**).

**Figure 4:**
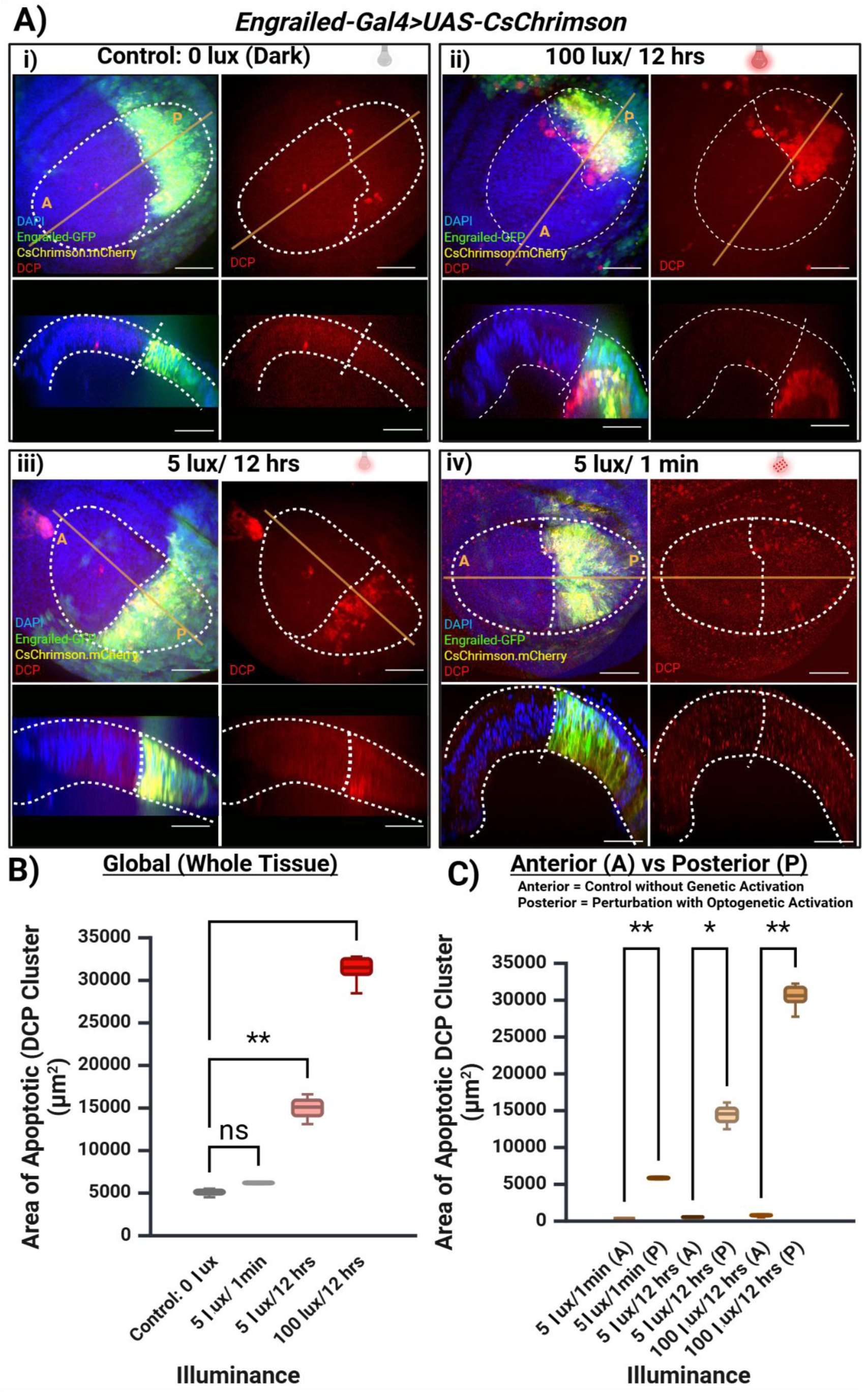
CsChrimson activation controls apoptosis. **A)** Max-Z projections and cross-sections of wing discs overexpressing CsChrimson with an Engrailed driver on the posterior compartment of the tissue and stained against apoptotic cells. Illuminance levels (100 lux or 5 lux) and period of activation (12 hr or 1 min) stated above the respective panels. Dotted white lines show the wing disc pouch (region of interest) and the division between Engrailed expression. Scale bar = 50 µm. **B)** Global quantification and comparison of apoptotic clusters across the whole wing disc. **C)** Compartment-specific quantification and comparison of apoptotic clusters in the anterior (A, non-perturbed) vs posterior (P, CsChrimson positive) compartments. n =7 for all cases. For statistics, Kruskal-Wallis test with Dunn’s multiple comparisons test was performed. P-values correspond to stars as described in Figure 2.

We first quantified apoptotic clusters in a negative control group: *CsChrimson* overexpressed in the posterior compartment using *Engrailed-Gal4*, reared in darkness and dissected under blue light without red-light activation (**Figure 4A-i**). This established baseline apoptosis levels in the wing disc. Upon continuous 100 lux red-light activation for five days until the dissection of third instar larvae, we observed a marked increase in apoptosis in the CsChrimson-expressing posterior half relative to both the anterior internal control compartment of the tissue and the 0 lux-grown control discs (**Figure 4A-i, ii, B, C**). These findings align with the severe adult phenotypes previously observed under 100 lux activation (**Figure 2B**). We also investigated optogenetic activation at 100 lux pulsed every 1 minute, but it resulted in insignificant change in apoptosis compared to 100 lux for 12 hours (**Figure S5).** To validate this effect across different spatial expression domains, we repeated the experiment using the *MS1096-Gal4* driver, which drives *CsChrimson* expression predominantly on the dorsal compartment of the wing disc **(Figure S1)**. A similar elevation in apoptosis was observed in the activated dorsal region (**Figure S6**). Together, these results demonstrate that excessive optogenetic stimulation leads to cytotoxicity locally, on the activated compartment of the tissue, triggering apoptosis in the developing tissue and contributing to deformities in the adult wing.

We observed significantly less apoptotic clusters within the *Engrailed-Gal4*–driven posterior compartment after 5 days of 5 lux red-light activation for 12 hours compared to the apoptosis levels at the 100 lux/12-hour condition. However, the apoptotic rate in the posterior region of the 5 lux/12 hrs condition remained elevated relative to the anterior (non-activated) compartment of the same disc (**Figure 4A-iii, B, C**). To further suppress apoptosis, we employed a 1-minute on/off duty cycle at 5 lux to target activation of the ERK signaling pathway as shown in the organ-related results. This stimulation pattern led to even less apoptotic activity. While the posterior compartment still showed slightly higher apoptosis than the anterior, the overall difference between apoptosis in discs in darkness and those activated with 5 lux/1-minute pulses was no longer statistically significant (**Figure 4A-i, iv, B, C**). Together, these results demonstrate that optogenetic activation of CsChrimson modulates apoptosis in a light intensity and duty cycle–dependent manner, with dimmer, pulsatile red light stimulation minimizing cell death compared to constant high-intensity exposure.

To determine whether these same activation patterns also influence cell proliferation, another closely related key process in tissue growth, we next examined mitotic activity in the developing wing disc. We quantified mitotic activity under the same optogenetic activation conditions used to assess apoptosis. *Engrailed-Gal4* driven *CsChrimson* overexpressed in *Drosophila* larvae reared entirely in darkness for 5 days and dissected under blue light, served as our baseline control for the number of mitotic cells in the wing disc pouch **(Figure 5A-i**). Under constant 100 lux red light exposure for 12 hours daily until dissection, overall mitotic activity in the wing pouch was slightly but significantly increased compared to the dark control (**Figure 5A-i, ii, B).** Moreover, no significant difference in mitotic cell count was observed between the posterior (CsChrimson-activated) and anterior (internal control) compartments (**Figure 5A-ii, C**).

**Figure 5:**
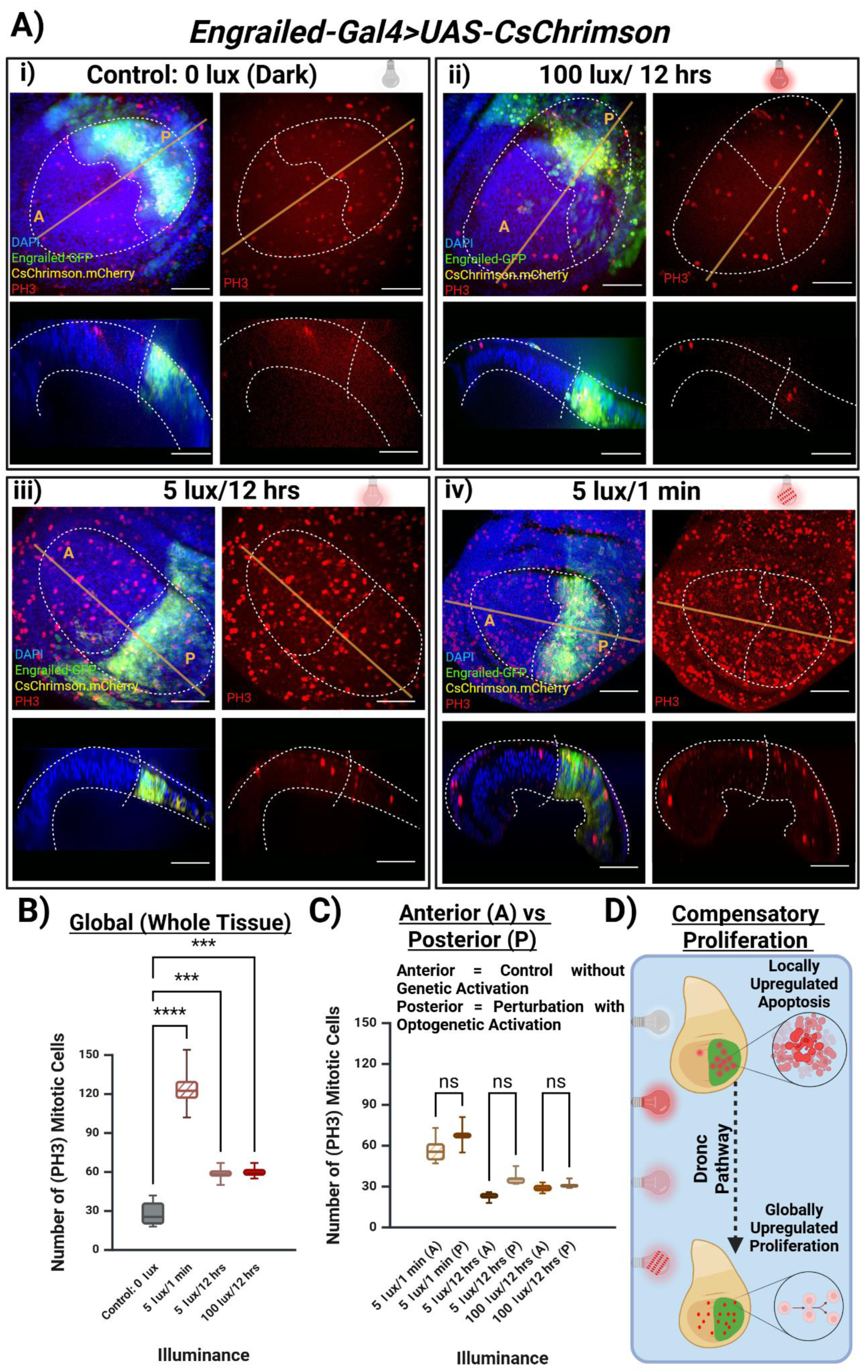
CsChrimson Activation Leads to Compensatory Proliferation. **A)** Max-Z projections and cross-sections of wing discs overexpressing CsChrimson on the posterior compartment of the tissue and stained against mitotic cells. Illuminance levels (100 lux or 5 lux) and period of activation (12 hr or 1 min) stated above the respective panels. Scale bar = 50 µm. **B)** Global quantification and comparison of the number of proliferating cells across the whole wing disc. **C)** Compartment-specific quantification in and comparison of the number of mitotic cells in the anterior (A, non-perturbed) vs posterior (P, CsChrimson positive) compartments. n > 5 for all cases. **D)** Pathway^54^ and schematic of apoptosis leading to compensatory proliferation in the wing disc. Kruskal-Wallis test with Dunn’s multiple comparisons test was performed. P-values correspond to stars as described in Figure 2.

When the stimulation pattern was changed to a 1-minute on/off duty cycle at 100 lux, we observed a modest, though statistically non-significant, increase in mitotic cells compared to dark conditions (**Figure S5**). Reducing light intensity to 5 lux while maintaining the same 12-hour activation period led to a significant increase in overall mitosis relative to the dark control (**Figure 5A-i, iii, B**). However, this increase was, again, not compartment-specific and mitotic rates remained similar between the posterior and anterior compartments despite optogenetic activation being restricted to the posterior, indicating a global effect across the tissue as opposed to locally spatial effect based on genetic perturbation.

Interestingly, the combinations of illumination intensities and activation periods showed a clear and opposing effect on apoptosis and proliferation within the wing imaginal disc. While apoptosis levels varied significantly depending on the specific light activation parameters, mitotic activity consistently increased across all light-exposed conditions compared to the dark-grown control. Notably, this increase in mitosis occurred even when apoptosis remained elevated, suggesting a compensatory mechanism at play (**Figure 5D**). This pattern aligns with the phenomenon of compensatory cell proliferation, wherein apoptotic cell death in one region of a developing tissue induces proliferation in neighboring cells to maintain organ size and tissue integrity^54–56^. In *Drosophila*, this mechanism is mediated by distinct molecular pathways depending on the developmental state of the tissue. In proliferating tissues such as the wing imaginal disc, the initiator caspase Dronc activates the Jun N-terminal kinase and p53 pathways, which in turn induce mitogenic signals like Decapentaplegic and Wingless to stimulate compensatory proliferation. In more differentiated tissues, effector caspases like DrICE and DCP-1 can activate alternative pathways, such as Hedgehog signaling, to drive a similar proliferative response^54^. This dual response of elevated apoptosis alongside increased mitosis under specific light conditions was consistently observed in our optogenetic experiments, reinforcing a role for compensatory proliferation and coherently linking the modulation of apoptosis to growth dynamics in the developing wing disc.

Building on our findings that CsChrimson-mediated optogenetic activation can fine-tune organ size through precise control of apoptosis and proliferation, we explored whether this tool could also be applied to neoplastic and cancer-like tissue states. In the robustly developing wing disc, we showed that specific light activation conditions promoted optimum organ growth, while excessive activation triggered apoptosis and tissue integrity loss, demonstrating the bidirectional control of tissue dynamics. These results prompted us to investigate whether tuning optogenetic activation could similarly influence dysregulated growth in a *Drosophila* neoplastic wing disc tumor model. Specifically, we asked whether adjusting the amplitude and timing of CsChrimson activation could reduce tumor burden or induce remission in neoplastic tissues, offering insights into how optogenetics might be used to control tumor growth through endogenous cellular signaling pathways.

### Light-Induced Modulation of Tumor Growth Through CsChrimson and Oncogenic Ras^V12^ Co-expression

Tumor growth is driven by complex and often dysregulated signaling pathways that are challenging to manipulate with spatial and temporal specificity^57^. To address this, we developed an optogenetic approach by co-expressing *CsChrimson* with the oncogenic *Ras^V12^* under the control of the *Patched-Gal4* driver, which expressed only along 5 cell widths at the anterior-posterior boundary, allowing tumor growth in a smaller region of the tissue while the rest of the tissue can survive without oncogenic expression. This enables precise light-mediated modulation of tumorigenic signaling *in vivo.* This strategy enables real-time, light-controlled investigation of oncogene-driven neoplastic growth, offering a novel platform for studying tumor dynamics and potentially informing light-based therapeutic interventions.

As a baseline, we first characterized the *Patched-Gal4* driver line, which exhibits a narrow and defined expression pattern along the anterior-posterior (AP) boundary of the wing disc^58^ (**Figure 6A, S1**). In the absence of *Ras^V12^* and *CsChrimson* expression, this line displayed normal wing disc morphology and served as a reference for spatial patterning and tissue shape (**Figure 6A i–iii**). When *Ras^V12^* and *CsChrimson* were co-expressed and flies were reared in darkness, CsChrimson remained inactive, and *Ras^V12^* expression alone induced pronounced tumorous overgrowth along the AP boundary, as visualized by GFP-tagged *Ras^V12^* and mCherry-tagged *CsChrimson* (**Figure 6B i–iii**). The wing disc morphology became distorted, reflecting unregulated neoplastic expansion along Patched expression pattern.

**Figure 6:**
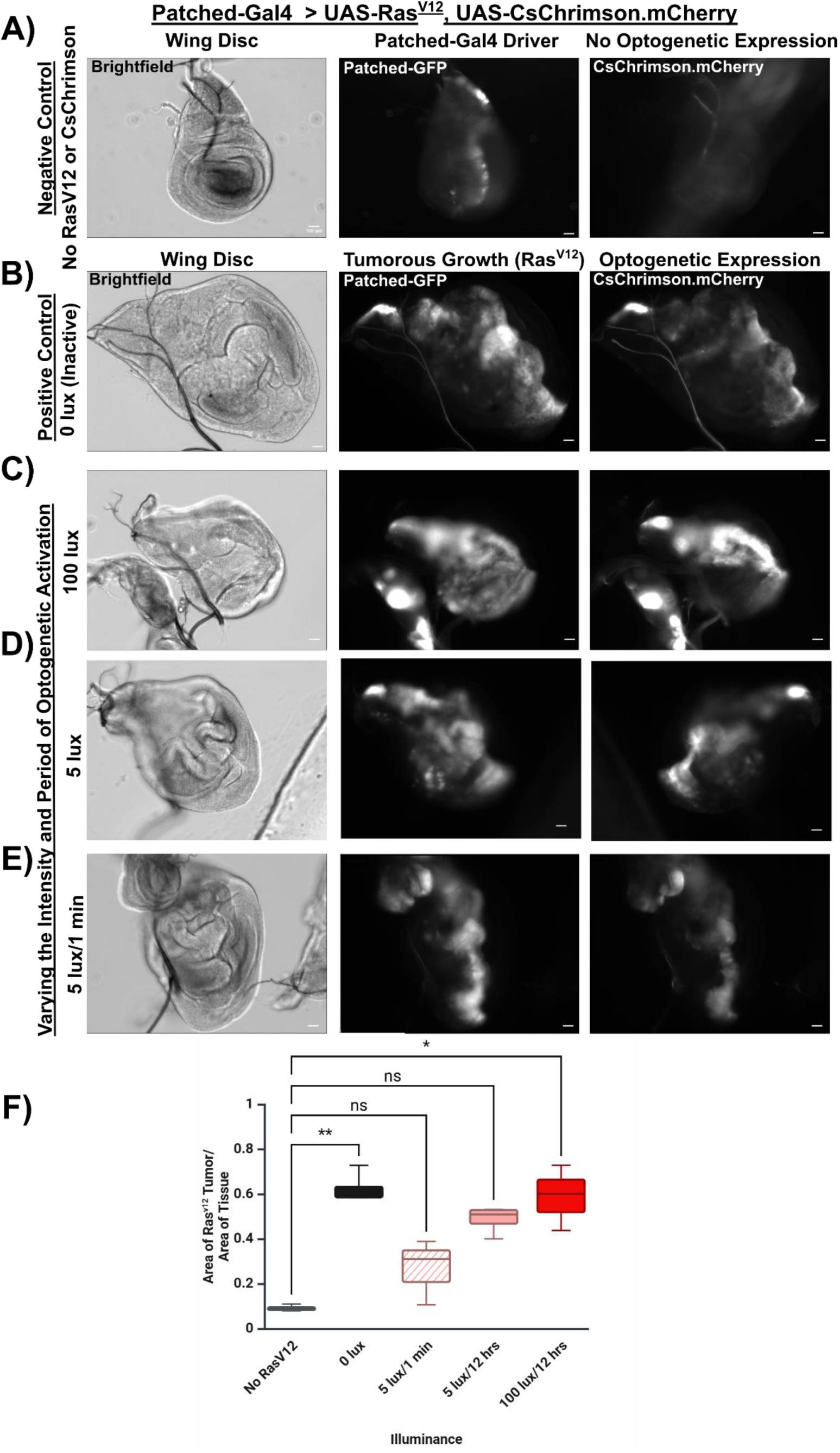
Systematic Optogenetic Activation Tunes Ras^V12^-mediated Tumorous Growth. Patched-Gal4 was used to drive co-expression of CsChrimson and Ras^V12^. **A)** (i) Control wing disc without tumor and (ii) Patched expression pattern without any (iii) CsChrimson and Ras^V12^ expression. **B) - E)** Gradual tuning of optoegentic activation of CsChrimson co-expressed with tumours Ras^V12^ gene showing (i) morphological shape changes of wing disc, (ii) changes is Patched pattern mediated by Ras^V12^, and (iii) CsChrimson co-expression confirmation for B) 0 lux, C) 100 lux, D) 5 lux, and E) 5 lux with 1 min On-Off duty cycle. Red arrows highlight changes in tumor shape. Scale bar = 100 µm. n > 3 for all cases. **F)** Boxplot quantification of tumorous areas over the entire wing disc areas for different illumination cases. Kruskal-Wallis test with Dunn’s multiple comparisons test was performed. P-values correspond to stars as described in Figure 2.

Next, we exposed developing larvae to continuous 100 lux red light for 12 hours per day throughout larval growth. This strong optogenetic activation did not reduce tumor size; instead, the wing disc remained distorted, and the tumorous region persisted (**Figure 6C i–i**ii). However, when light intensity was reduced to 5 lux under the same activation period, we observed a surprising reduction in tumor size despite ongoing *Ras^V12^* expression. Although overall disc morphology was still aberrant, the cancerous tissue marked by *Patched* expression showed clear signs of regression (**Figure 6D i-iii)**. Most notably, implementing a 5 lux red light pulsed in a 1-minute on/off duty cycle for five days led to even greater tumor remission within the *Patched*-driven domain (**Figure 6E i-iii)**. Quantification of the tumor area shows a non-significant difference between the negative control case and both the 5 lux/12 hrs and 5 lux/1 min cases (**Figure 6F**).

These results demonstrate that tuning optogenetic activation through both intensity and temporal dynamics can significantly influence tumor overgrowth in a tissue. Quantification of the tumor size reflects a similar trend as wing size modulation (**Figure 6, 3).** Moderate and periodic activation of CsChrimson appears to induce cellular signaling environments that oppose oncogene-driven overgrowth, highlighting the potential for optogenetic tools to reshape the trajectory of cancer-like growth in a genetically tractable *in vivo* model.

## Discussion

This study demonstrates the potential of optogenetics as a tool for probing and manipulating developmental and pathological processes *in vivo* with spatiotemporal precision, specifically within the *Drosophila melanogaster* wing imaginal disc. By using the red-light-activated CsChrimson channel, we established that the spatiotemporal dynamics of calcium signaling directly influence cellular processes such as proliferation, apoptosis, and overall tissue morphogenesis **(Figure 1)**. Our computational model provided deeper systems-level insight into the observed increase in Ca²⁺ dynamics upon stimulation. Emerging variants of GCaMP, with enhanced brightness and a broader range of temporal kinetics^59^, offer promising tools for more accurate quantification of Ca²⁺ activity, which could significantly improve the parameterization and predictive power of future computational models. Through fine-tuning the intensity and duration of light exposure duty cycles, we demonstrated a nuanced control over organ size, revealing the importance of intracellular calcium fluxes in developmental regulation (**Figure 2, 3)**. These findings suggest the presence of a “Goldilocks zone”^60^, an optimal level of cytosolic Ca²⁺ and cation transport through the optogenetic channel that promotes robust organ growth.

We observed that higher light intensity (100 lux) led to increased apoptotic cell death and a concurrent decrease in cell proliferation, particularly within the optogenetically perturbed compartments. In contrast, reduced light intensity (5 lux) or intermittent light activation (1 min on/off cycle) resulted in a striking increase in mitosis accompanied by lower apoptosis levels. These findings highlight a delicate balance between apoptosis and proliferation that is critical for proper organ development and align with studies showing that patterned apoptosis contributes to local growth regulation and final tissue shape during wing morphogenesis^61^. This also aligns with existing literature showing that patterned apoptosis actively contributes to regulating local growth and final tissue shape, highlighting cell death as a key modulator of organ size alongside proliferation during wing morphogenesis^61^. While these results suggest that optogenetically induced apoptosis, especially under high or prolonged activation, may trigger a non-autonomous increase in mitosis via compensatory proliferation, contributing to the enlarged wing phenotype observed under specific light conditions (**Figure 3, 4, 5**), this interpretation remains preliminary and warrants further investigation into signaling pathways in future studies.

Interestingly, we also observed a light-induced non-cell-autonomous effect, where tissue-wide proliferation increased, even in compartments not directly subjected to light activation. This effect aligns with the compensatory proliferation mechanism previously described in *Drosophila* tissues, where apoptosis triggers mitogenic signals to promote tissue regeneration^55^ (**Figure 5**). This finding is particularly compelling, as it suggests that light-activated signaling in one compartment can induce a non-autonomous, global increase in proliferation throughout the tissue. Together, these results demonstrate that optogenetic modulation of signaling not only alters cell fate locally but can also propagate system-wide changes in proliferation dynamics. Our data further emphasizes that calcium dynamics are central to coordinating these cellular responses during development.

The application of optogenetics to a *Ras^V12^*-driven tumorigenic model revealed an added dimension to this approach (**Figure 6)**. By adjusting the intensity and pattern of CsChrimson activation, we successfully influenced the growth and regression of *Ras^V12^*-induced tumors in wing discs. Tumors persisted under high-intensity light but shrank significantly when activation was set to 5 lux and further reduced with pulsed 5 lux light conditions. Additionally, our results showed that loss of integrin function resulted in disrupted epithelial organization and enhanced invasive behavior within the wing disc epithelium at prolonged optogenetic activation, consistent with the established role of integrins in maintaining tissue architecture; such dysregulation may be investigated in the *Ras^V12^* to investigate cell detachment and migration, key early steps in metastatic progression^62^ (**Figure 2, 6).** These results and insights suggest that optogenetic modulation of growth-related signaling cascades could offer new strategies for tumor control, opening up a potential therapeutic avenue for manipulating neoplastic growth.

Our findings align with existing literature demonstrating the importance of ion-transporting channels and calcium signaling in regulating critical cellular functions, such as apoptosis, proliferation, and differentiation^63–66^. In *Drosophila* wing discs specifically, prior studies have shown that mechanical stress or genetic perturbations can trigger spatially patterned Ca^2+^ waves that influence morphogenesis^67^. Our observation that different patterns and intensities of optogenetically induced Ca^2+^ activity lead to opposing effects on apoptosis and proliferation is consistent with known roles of Ca^2+^ in regulating both pro-survival and pro-death pathways^68^. The compensatory proliferation we observed in response to increased apoptosis mirrors previous work identifying Dronc and effector caspases as key mediators linking apoptotic signaling to mitogen induction through pathways such as JNK and Hedgehog^54^. Furthermore, our tumor model using *Ras*^V12^ driven by the *Patched-Gal4* driver echoes the cancerous transformation observed in other *Drosophila* models of oncogenic *Ras* signaling^69^, and our ability to modulate tumor burden with graded optogenetic activation suggests that calcium-modulated growth-related signaling cascade may be harnessed to fine-tune growth or induce remission. These connections reinforce the relevance of our optogenetic platform not only for basic developmental biology but also as a conceptual framework for therapeutic intervention in cancer.

In conclusion, this study highlights the potential of optogenetic control of calcium signaling and growth morphogenetic processes as a powerful tool for regulating organ development, tissue homeostasis, and even tumor growth. Our findings open new possibilities for using precise light-based interventions to study and treat developmental disorders and abnormal tissue growth, offering a promising direction for future therapeutic applications.

### Limitations of the Study

This study focuses on reporting key observations relating optogenetic control of physiological signaling on both wild-type and tumor growth, but future efforts are required to test rigorously inferred mechanisms of compensatory proliferation. Refinement of optogenetic tools will also be crucial for expanding the precision of these experiments. The use of Ca^2+^-specific channelrhodopsins like CapChR, which offer improved sensitivity and selectivity, could enable more precise control of calcium signaling with reduced off-target effects, enhancing the ability to finely tune cellular responses with lower light intensities^70^. Other ions transporting optogenetic tools, such as NaChRs^71^, kalium channelrhodopsins^72^, Cl-out^73^, may also provide precise temporal and spatial control over specific ion fluxes, enabling detailed studies of cellular excitability, signaling pathways but their adaptability in *Drosophila* epithelial system is a limiting factor. This could provide more detailed control of tissue morphogenesis and offer better therapeutic strategies for diseases like cancer, where calcium dysregulation plays a significant role.

## Acknowledgments

This work is based upon efforts supported by the EMBRIO Institute, contract #2120200, a National Science Foundation (NSF) Biology Integration Institute. MSM and JZ were supported in part by the NIH Grant R35GM124935 and R35GM156615. SC was partially supported by NSF CBET-1941596. We thank Dr. Sara Cole for helping us use the confocal microscopes at the Optical Microscopy Core of Notre Dame Integrated Imaging Facility. The authors would also like to thank Dr. Maria Unger, Dr. Pablo Cisternas Esguep, Sayandeepa Raha, Dr. Nilay Kumar, Dr. Vijay Velagala, David Gazzo, Benjamin Speybroeck, Beatriz Borrelly, Dr. Daniel Laky and Molly Dougher for their insight and helpful discussions. BioRender.com was used to create figures and plots.

## Author Contributions

MSM conceptualized, designed and performed all experiments, analyzed and interpreted the data, prepared the manuscript, created figures, and developed quantification protocols. SC contributed to writing, figure creation and editing, ran model simulations, analyzed and interpreted the data, model and coding implementation. SC and AD designed and implemented the optogenetic calcium dynamics model. CF contributed to writing, figure editing and finalizing of the manuscript. ZW designed a genetic scheme for and created CsChrimson and Ras^V12^ co-expressed fly lines. JZ conceived, designed, analyzed, and interpreted results for all experiments and simulations, supervised this study, and contributed to writing the manuscript. All authors read and agreed to the publication of the manuscript.

## STAR Methods

### RESOURCE AVAILABILITY

#### Lead contact

Further information and requests for resources and reagents should be directed to and will be fulfilled by the lead contact, Jeremiah Zartman.

#### Materials availability

This study did not generate new unique reagents.

#### Data Availability

Data reported in this paper will be shared by the lead contact upon request.

#### Materials and Methods

### EXPERIMENTAL MODEL AND SUBJECT DETAILS

#### Fly stocks and culture

*Drosophila melanogaster* were reared at 25 °C under a 12-hour light/dark cycle or in complete darkness or under red light condition as stated in results. Virgin females were collected twice daily from Gal4 driver line cornmeal food bottles and crossed with males carrying the indicated UAS-transgene constructs (gene of interest) at a 15:5 female-to-male ratio in 1 mM all-trans Retinal supplemented cornmeal food. Crosses were staged for 8 hours to collect appropriately aged third instar larvae. Wandering third instar larvae of both sexes were dissected for wing imaginal disc excision. Adult flies were collected upon eclosion for wing morphology analysis. F1 larvae and pupae were sorted by genotype based on the crossing scheme to ensure correct expression. Adult progeny were collected 24 hours post-eclosion, separated by sex, and used for wing quantification.

#### Live Imaging to Study Calcium Dynamics

A pre-established wing disc imaging protocol was followed^74^. Wing imaginal discs were dissected under blue light, to avoid activation or desensitization of the CsChrimson channels before imaging, from physiological third instar crawling larvae on day 5 after egg laying (AEL) and immediately mounted into polyethylene terephthalate laminate (PETL) microfluidic devices for live imaging under continuous media flow^75^. The culture medium consisted of a supplemented Grace’s cocktail^76^, with Grace’s insect medium prepared without sodium bicarbonate and supplemented with 5 mM BisTris. The pH was adjusted to 6.6–6.7 at room temperature, and the media was stored at 4°C for up to one month. On the day of use, 5% FBS, 1× Penicillin-Streptomycin were added to prevent microbial contamination and 10 nm 20-Hydroxyecdysone was added to activate calcium signaling pathway. Wing discs were incubated in culture media that was supplemented with 100 µM all-trans Retinal for 15 minutes before imaging. The initial time points under darkness served as control discs prior to red light treatment. Imaging was performed using a custom Nikon/Andor spinning disc confocal microscope. Full z-stacks (∼120 images) were acquired every minute over 30 minutes using 40× magnification using an oil objective. Image stacks were processed in Fiji^77^, with brightness and contrast adjusted for visualization.

#### Calcium Dynamics Modeling

Computational models developed by Soundarrajan et al^17^ and Politi et al^78^ were adapted to emulate the optogenetic activation of the Ca^2+^ dynamics led by CsChrimson. This computational framework serves as a foundational model for simulating Ca^2+^ signaling dynamics within epithelial cells of the *Drosophila* wing disc, integrating both endogenous and optogenetically induced signaling pathways. Originally developed to capture intracellular inositol trisphosphate, IP_3_R-mediated calcium activity, the model represents core processes of Ca^2+^ homeostasis and signal propagation. It accounts for the generation and degradation of IP_3_, calcium release from the endoplasmic reticulum (ER) through IP_3_R, and active calcium reuptake into the ER via SERCA.

The calcium dynamics model was implemented in Python 3.8.19, using matplotlib 3.7.3, numpy 1.24.3, seaborn 0.12.2, and scipy 1.10.1 into a class object with defined functions for simulation and visualization^79–83^. A fixed geometry was used containing 860 cells, and the simulation time was set for 3600 seconds (1 hour) by default with a step size of 0.2 seconds, leading to a total number of 18000 finite elements per simulation. On a 32 GB RAM machine with Intel(R) Core(TM) i7-1255U processor, a simulation was performed in approximately 40 seconds, and each visualization was generated in approximately 6 seconds. The primary capabilities of the model are simulation and visualization. The original and optogenetic models are defined in equations 1-4 and 5-6, respectively and parameters for the model are stated in Table 1.

#### Model Equations form Soundarrajan et al

IP_3_ Dynamics

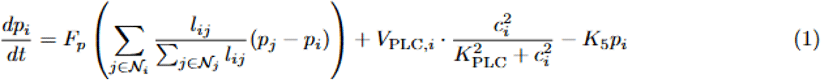

Ca^2+^ Dynamics

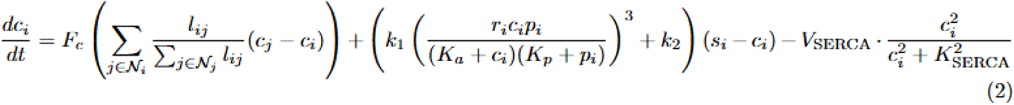

ER Calcium (Algebraic Expression)

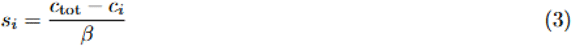

IP_3_R Inactivation Dynamics

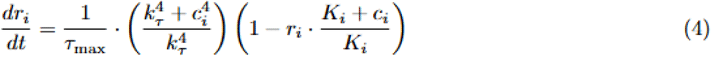

Optogenetic Extension

We incorporate optogenetic stimulation as an external calcium input term:

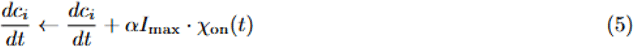

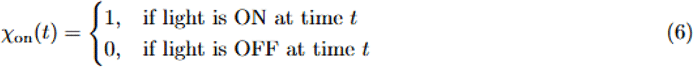

Parameter Table

**Table 1.**
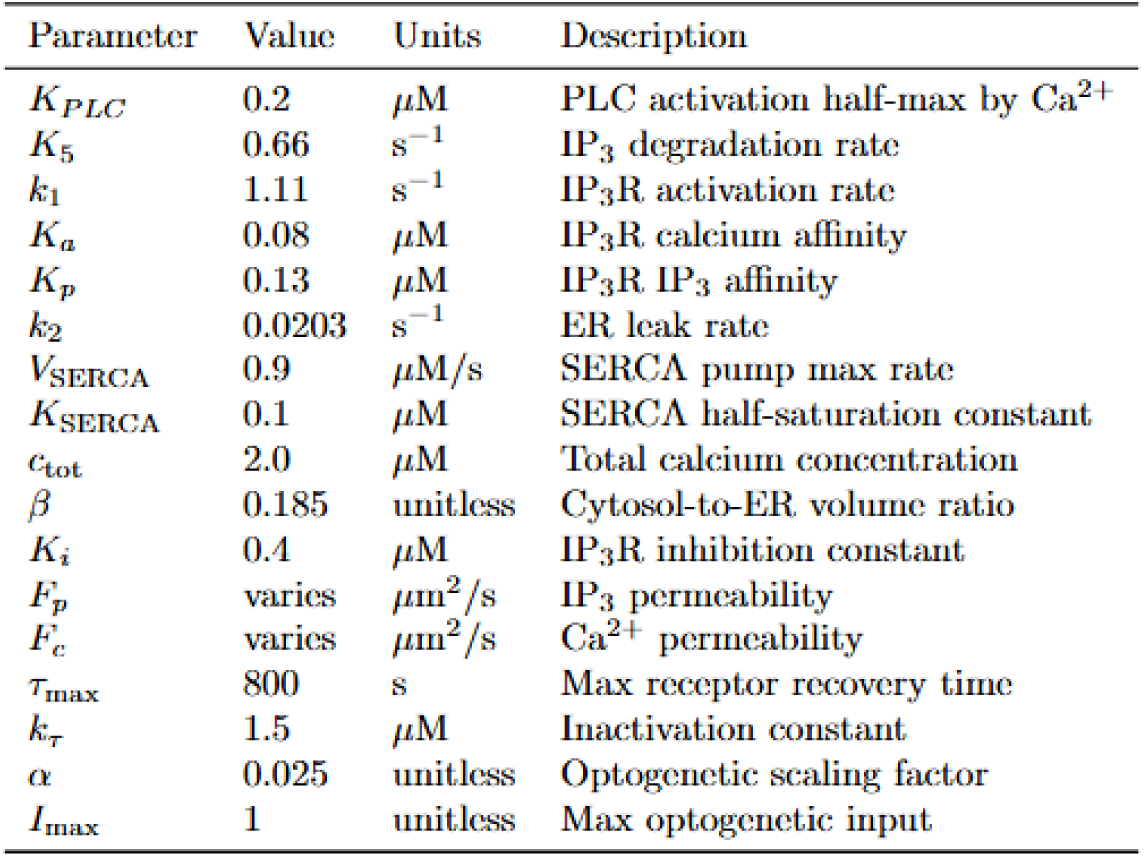
Model parameters based on Soundarrajan et al. for calcium and IP_3_ signaling with optogenetic extension.

Further details regarding the model are included in the Supplementary Information.

#### Wing Disc Immunohistochemistry and Mounting

Wing imaginal discs were dissected from third instar *Drosophila* larvae at 20-minute intervals under blue light to avoid optogenetic activation. Discs were fixed in ice-cold 4% paraformaldehyde for 30 minutes in PCR tubes. Following fixation, discs were rinsed three times with fresh PBT (PBS containing 0.03% Triton X-100). Tubes were placed on a nutator at room temperature for 10 minutes, followed by an additional PBT rinse; this nutation/rinse cycle was repeated three times. After the final rinse, 250 μL of 5% normal goat serum (NGS) in PBS was added to each tube, and samples were incubated on a nutator for 30 minutes at room temperature. The NGS was then replaced with 250 μL of primary antibody solution prepared in 5% NGS, and the samples were incubated overnight at 4 °C. The next day, three quick PBT rinses were performed, followed by three 20-minute nutation/rinse cycles in PBT. After the final rinse, 250 μL of secondary antibody solution in 5% NGS was added, and tubes were nutated for 2 hours at room temperature. This was followed by another set of three quick rinses and three 20-minute nutation/rinse cycles in PBT. Discs were then incubated overnight at 4 °C for a final wash and mounted in Vectashield mounting medium for imaging.

#### Confocal Microscopy for Fixed and Live Tissue Imaging

Wing imaginal discs were imaged using a Nikon Eclipse Ti confocal microscope equipped with a Yokogawa spinning disc, a Nikon A1R-MP laser scanning confocal microscope, or a Leica Stellaris 8 DIVE point scanning confocal microscope. Image acquisition was performed using an iXon EM+ cooled CCD camera with IQ3 software (Andor Technology, South Windsor, CT) for the spinning disc system, NIS-Elements software for the A1R-MP system, and LAS X microcope software for the Leica Stellaris DIVE system. Z-stacks were collected throughout the full depth of the disc with step sizes of 1 μm using 40× and 60× oil immersion objectives. Imaging parameters included a 200 ms exposure time and 44% laser intensity using 405 nm, 488 nm, 561 nm, and 640 nm lasers at 50 nW power. Scanning was performed from the apical to basal surface, beginning with peripodial cells followed by the underlying columnar epithelium. Optical slices were acquired at intervals equal to half the compartment length to ensure appropriate spatial resolution.

In Figure 4 and 5, we manually assessed the ratio of antibody-stained cells to nuclei (DAPI) co-stained to quantify apoptotic and mitotic cells labeled with *Drosophila* caspase-1 (DCP-1)^53^ and Phospho-Histone H3 (PH3)^84^ antibodies, respectively. By comparing these counts, we derived the apoptotic area and the number of mitotic cells within the wing disc pouch area.

#### Optogenetic Experimental Setup

To ensure keeping the optogenetic channel inactivated until intended, the flies grown in darkness were dissected under blue (∼400 nm wavelength) light, as this wavelength does not activate the CsChrimson channel. As seen in the schematic, an optogenetic experimental setup for *Drosophila* requires multiple steps (**Figure S2**). First, a prerequisite for the functional generation of CsChrimson in *Drosophila* is the supplementation of 1 mM All-Trans-Retinal. To ensure optimal conditions for optical activation, the crosses were set up in darkness since white light can desensitize and deactivate the optogenetic channels. The dark condition was maintained by wrapping the vials in aluminum foil. On collecting the embryos in darkness, light exposure of the channelrhodopsin-specific wavelength of light, which is red light (∼600 nm) for CsChrimson, was initiated using 1.5 watts LED strips at the 1st instar larval stage in aluminum foil-wrapped 6 x 4 x 6 inch paper boxes. The larvae were left under this monochromatic light condition until they pupariated and the vial moved back to darkness until eclosion. 12 hours of darkness and 12 hours of illuminated conditions were maintained to refrain from perturbing the circadian rhythm. At the 3rd instar crawling larval stage, the wing disc can be dissected for Ca^2+^ sensing and imaging. Otherwise, to observe the adult wing phenotype, the flies were kept in darkness until the emergence of F1 adults. All flies were grown at 25° C. An Arduino microcontroller appended with transistors for amplification and resistors for dimming light was powered by a 9 V DC battery and programmed to maintain the period of illuminance (**Figure S3**). A digital illuminance meter was used to measure the level of light in lux (Digital Light Meter, LX1330B, Dr. Meter). A conversion from illuminance in lux to irradiance in mW/m^2^, along with the values of the resistors used for each illumination level.

#### Statistical analysis

Calcium spiking analysis in Figure 1D, and analysis of wing area variation for 5 lux/ 12 hrs vs 5 lux/ 1min illuminance in Figures 3D and 3F was done using Unpaired t-test with Welch’s correction^85^. Analysis of wing area variation for varying illuminance in Figure 3C and 3E, and analysis for global and compartment-specific cell death and cell proliferation levels in Figures 4B, 4C, 5B and 5C was done using Kruskal-Wallis test with Dunn’s multiple comparisons test^86,87^. All statistical analysis data are included in the Supplementary Information as Tables S2-S11.

## Supplemental Item Titles

1. **Model description and computational details**. Related to Figure 1.
2. **Figure S1**: Wing imaginal disc of *Drosophila melanogaster* as a testbed for precise gene expression. Related to Figures 1-6.
3. **Figure S2**: Schematic of optogenetic experimental setup. Related to Figures 1-6.
4. **Figure S3**: Schematic of the microcontroller-based electrical circuit. Related to Figures 1-6.
5. **Figure S4**: Computational model provides further insights into CsChrimson-mediated Ca2+ Dynamics. Related to Figure 1.
6. **Figure S5**: Apoptosis and Proliferation at 100 lux/1 min illumination. Related to Figures 4 and 5.
7. **Figure S6**: Apoptosis and mitosis levels on optogenetic activation. Related to Figures 4 and 5.
8. **Table S1**: Conversion from Illuminance (lux) to Irradiance (W/m^2^).
9. **Table S2**: Calcium spiking statistical analysis using Unpaired t-test with Welch’s correction. Related to Figure 1D.
10. **Table S3**: Statistical analysis of female wing area variation for varying illuminance using Kruskal-Wallis test with Dunn’s multiple comparisons test. Related to Figure 3C.
11. **Table S4**: Statistical analysis of male wing area variation for 5 lux/ 12 hrs vs 5 lux/ 1min illuminance using Unpaired t-test. Related to Figure 3D.
12. **Table S5**: Statistical analysis of female wing area variation for varying illuminance using Kruskal-Wallis test with Dunn’s multiple comparisons test. Related to Figure 3E.
13. **Table S6**: Statistical analysis of female wing area variation for 5 lux/ 12 hrs vs 5 lux/ 1min illuminance using Unpaired t-test. Related to Figure 3F.
14. **Table S7**: Statistical analysis of global (tissue wide) apoptosis for varying light illuminance levels using Kruskal-Wallis test with Dunn’s multiple comparisons test. Related to Figure 4B.
15. **Table S8**: Statistical analysis of compartment-specific (anterior vs posterior) apoptosis for varying light illuminance levels using Kruskal-Wallis test with Dunn’s multiple comparisons test. Related to Figure 4C.
16. **Table S9**: Statistical analysis of global (tissue wide) mitosis for varying light illuminance levels using Kruskal-Wallis test with Dunn’s multiple comparisons test. Related to Figure 5B.
17. **Table S10**: Statistical analysis of compartment-specific (anterior vs posterior) mitosis for varying light illuminance levels using Kruskal-Wallis test with Dunn’s multiple comparisons test. Related to Figure 5C.
18. **Table S11:** Statistical analysis of the area of tumor over the area of whole wing disc using Kruskal-Wallis test with Dunn’s multiple comparisons test. Related to Figure 6F.

