## Supplementary Information for "Forward Engineering Organ Development and Cancer Therapeutics with Optogenetics"

18. **Table S11:** Statistical analysis of area tumor over area of whole wing disc using Kruskal-Wallis test with Dunn's multiple comparisons test. Related to Figure 6F.

**Model description and computational methods:**

To integrate optogenetics into the original referenced model[1], the term $u_i$ (t) was introduced. This term, described in the Methods section of the manuscript, includes α, $I_{max}$, and X(t), where X(t) is a step function indicating the activation and deactivation of the optogenetic channel via red light. For this work, the model assumptions dictate that red light is always on, unless labeled dark, for all simulations. Tissue geometry is fixed with 860 cells. Tissue generated using Voronoi tessellation, followed by Lloyd's relaxation to make shapes more uniform[2]. The geometry is described computationally using Laplacian, adjacency, and vertex matrices, contributing to the model's diffusion terms. The fraction of initiator cells was set to $1/(\text{number of cells})^{0.8}$ as defined for the original model. No parameters related to tissue damage or apoptosis were implemented, and control using derivatives is not meant to imply the exact mechanism of behavior with optogenetic use of CsChrimson. As described in the main text, the gap junctions are disabled for the main simulations due to the optogenetic light increasing the value of α and decreasing the diffusion coefficients.

All coding and simulations were completed in Python, using standard libraries and packages. The model derivatives were simulated using a manual forward Euler approach in a for loop for each step. 18000 total steps were utilized, with a simulation time of 3600 seconds and 0.2-second step size due to the model's stiffness. Simulation time was varied by modifying the total model time for both the simulation and stimulus. Graphics were developed using embedded functions within the "Pouch" class object, utilizing seaborn and matplotlib for customized visualizations, including $V_{PLC}$ profile, and max calcium profile.

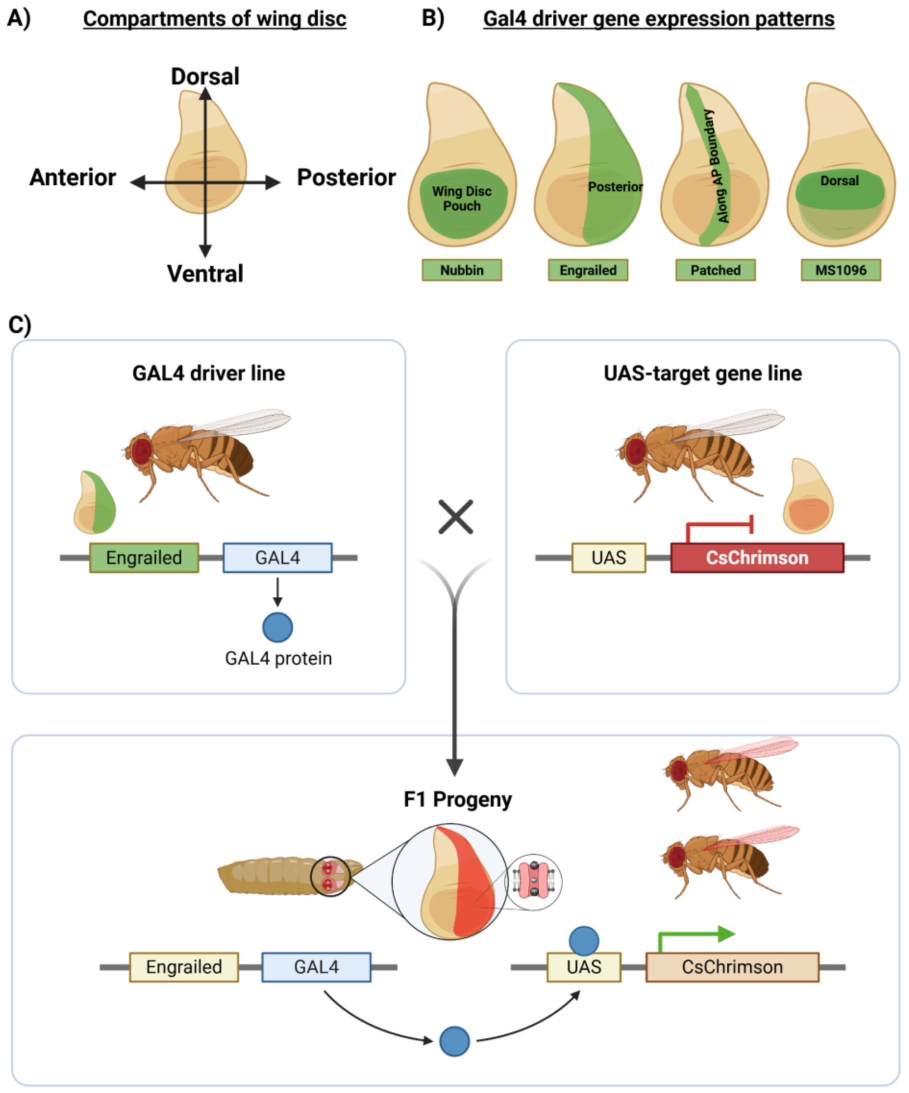


**Figure S1: Wing imaginal disc of *Drosophila melanogaster* as a testbed for precise gene expression. Related to Figures 1-6. A)** The four compartments of the wing disc: anterior (A), posterior (P), dorsal (D) and ventral (V) giving rise to A-P and D-V boundaries. **B)** Expression patterns of the Gal4 drivers used in this manuscript: *Nubbin*: in the entire wing disc pouch region, *Engrailed*: in the posterior compartment of the tissue; making the anterior compartment an internal control without genetic perturbation, Patched: Along 5 cell widths at the Anterior-Posterior boundary, and *MS1096*: predominantly on the dorsal compartment. **C)** UAS-Gal4 bipartite system driving the expression of a gene of interest, *CsChrimson*, in a spatial pattern dictated by the Gal4 driver of interest, which is *Engrailed* here..

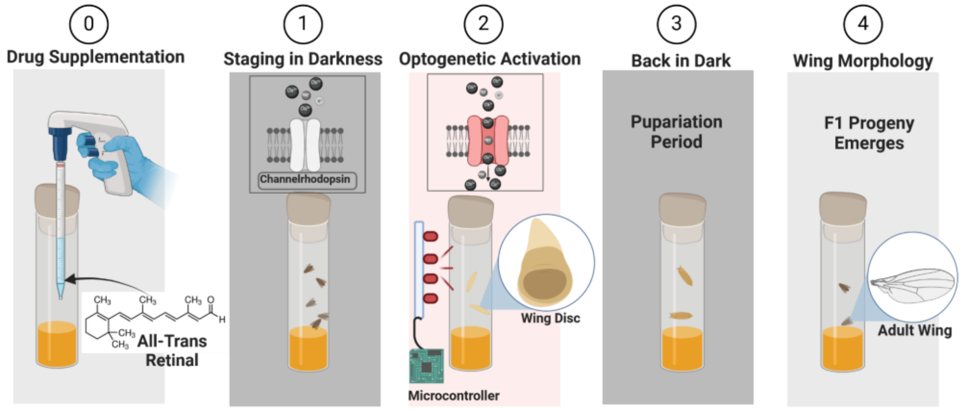


**Figure S2: Schematic of optogenetic experimental setup. Related to Figures 1-6.** Step-by-step guide to setting up an optogenetic experiment for *Drosophila* epithelial tissue include step 0: adding All-Trans Retinal to to the food, step 1: Wrapping the food vial in aluminum foil to protect from white light exposure, step 2: Using microcontroller-activated LED lights for systematic optogenetic activation, step 3: Put the food vial containing F1 progeny back in darkness once they pupariate, and step 4: Collect F1 progeny with optogenetic activation to evaluate terminal organ morphology.

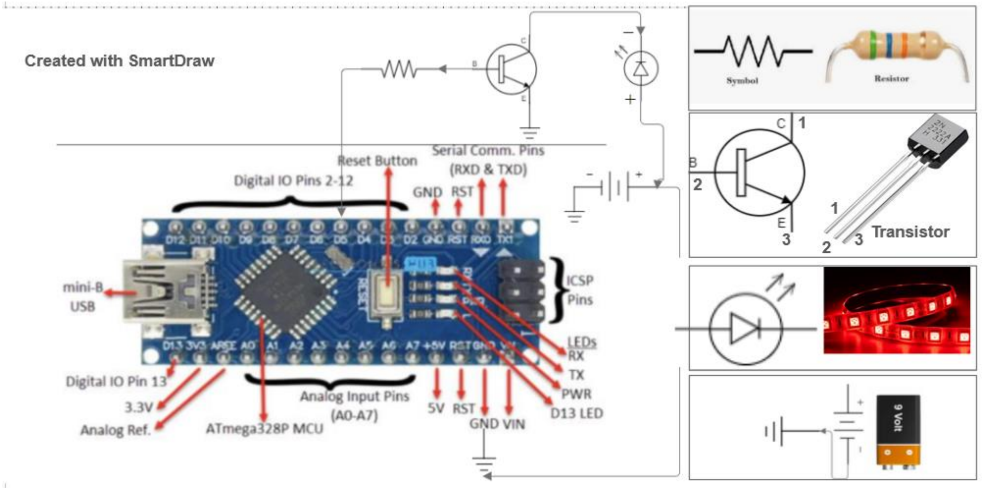


**Figure S3: Schematic of microcontroller-based electrical circuit. Related to Figures 1-6.** An Arduino Nano microcontroller is used to code and systematically vary the intensity and period of red light activation using a circuit designed with resistors and a transistor, powered by a 9 V battery.

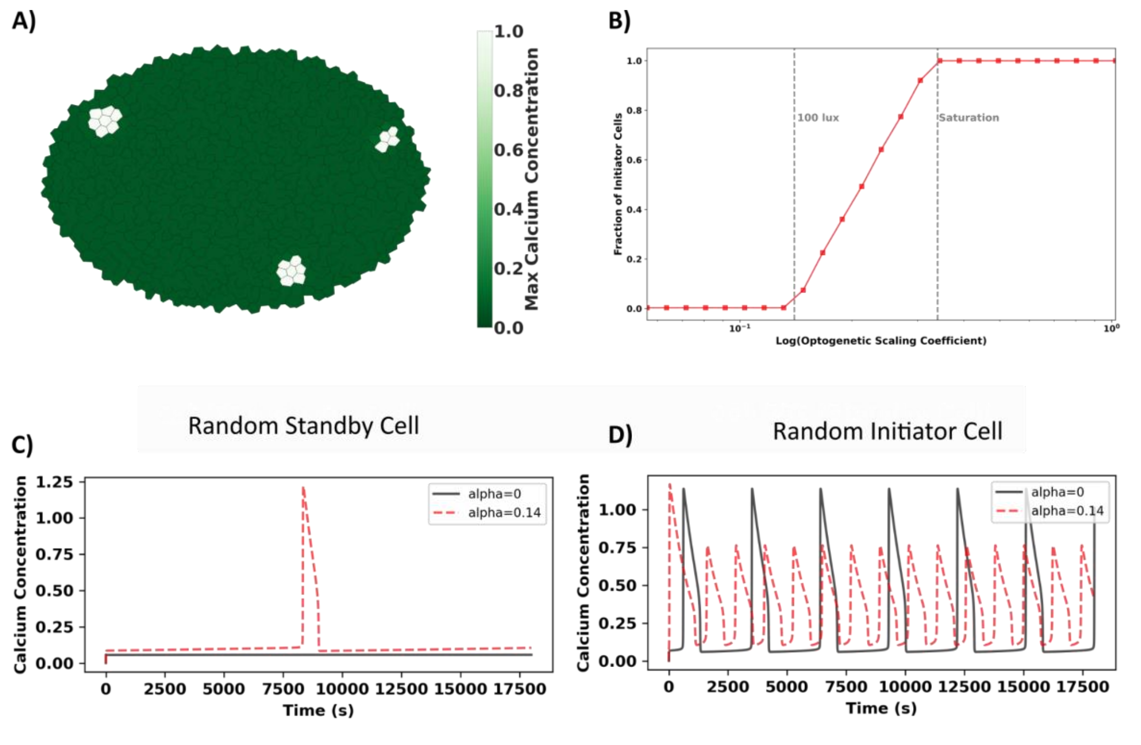


**Figure S4: Computational model provides further insights into CsChrimson-mediated $Ca^{2+}$ Dynamics. Related to Figure 1. A)** Simulated wing disc with open gap junction showing intercellular transient-like $Ca^{2+}$ dynamics (white cells) at 30 minutes, which was not observed on CsChrimson activation. **B)** Optogenetic scaling coefficient, α showing $Ca^{2+}$ activation and saturation of initiator cells. **C-D)** Randomly selected 860 total cells, C) a standby cell and D) an initiator cell showing the effect of varying optogenetic scaling coefficient, α at 0 and 0.14, as interpreted from panel B.

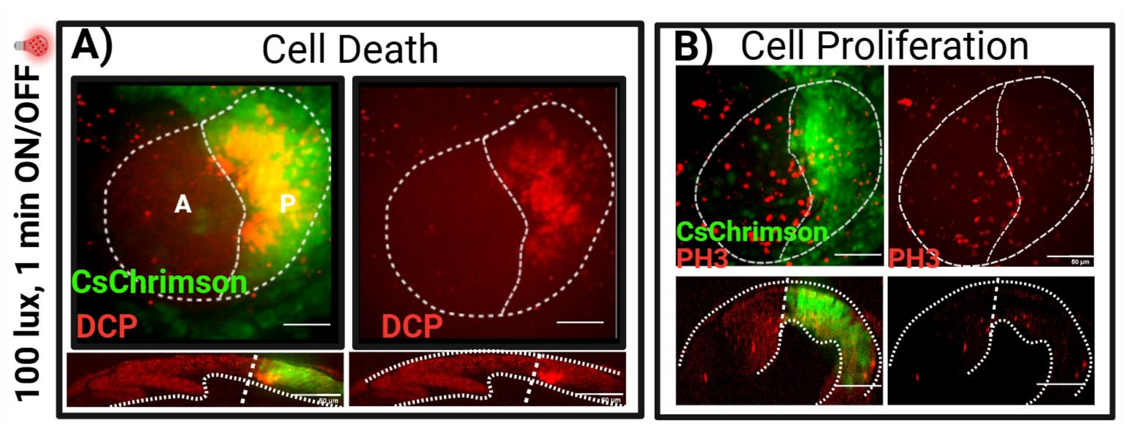


**Figure S5: Apoptosis and Proliferation at 100 lux/1 min illumination. Related to Figures 4 and 5. A) apoptosis B) proliferation**

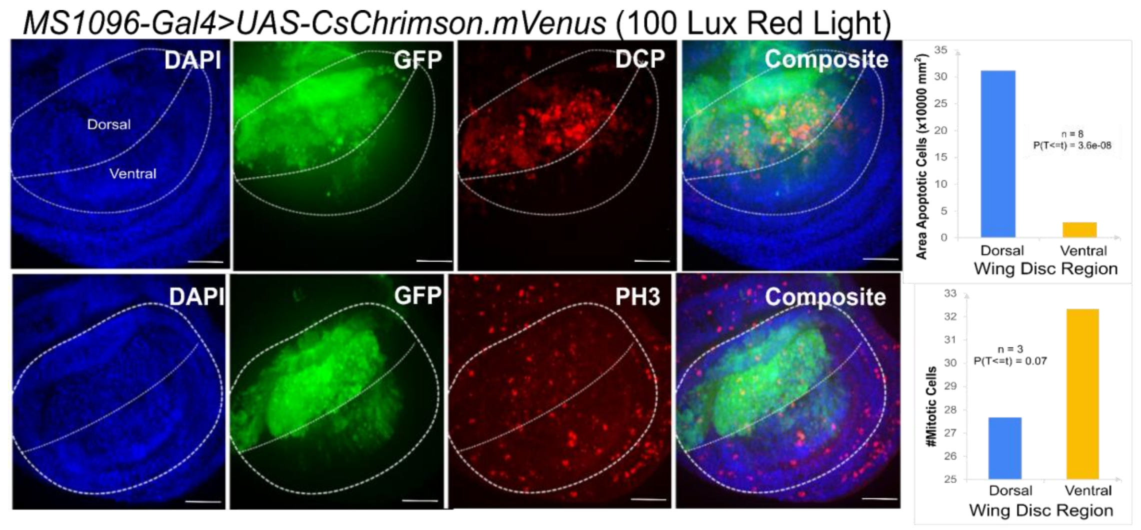


**Figure S6: Apoptosis and mitosis levels on optogenetic activation. Related to Figures 4 and 5.** 100 lux red light activation during 5 days of embryonic to larval growth in CsChrimson overexpressed predominantly on the dorsal compartment of the tissues driven by MS1096-Gal4 driver. On the top panels, we observe upregulation of cell death, stained by DCP, on the dorsal compartment as compared to the ventral compartment.. On the bottom panels, PH3 staining reveals no significant changes in proliferation levels on dorsal vs ventral compartments. Scale bar = 100 µm. n = 9.

**Table S1: Conversion from Illuminance (lux) to Irradiance (W/m$^2$). lux to watts calculation with area in square meters.** The power P in watts (W) is equal to the illuminance $E_v$ in lux times the surface area $A$ in square meters (m$^2$), divided by the luminous efficacy $\eta$ in lumens per watt (lm/W): $P_{(W)} = E_{v(lx)} \times A_{(m^2)} / \eta_{(lm/W)}$. For 600 nm red light, the conversion is as follows:

| **Illuminance (lux)** | **Irradiance (mW/m$^2$)** | **Resistance in Circuit (Ω)** |
|---|---|---|
| 0 | 0 | N/A |
| 3 | 6.96 | 4000 |
| 5 | 11.6 | 3000 |
| 15 | 34.8 | 1500 |
| 30 | 69.60 | 1000 |
| 50 | 116.02 | 750 |
| 70 | 162.44 | 470 |
| 80 | 185.86 | 200 |
| 90 | 209.28 | 75 |
| 100 | 232.71 | 0 |

**Table S2: Calcium spiking statistical analysis using Unpaired t-test with Welch's correction. Related to Figure 1C.**

| t | DF | p-value (two-tailed) | Difference Between Means (A - B) ± SEM | 95% Confidence Interval | Cohen's *d* |
|---|---|---|---|---|---|
| -27.7128 | 7.3529 | <0.0001 **** | 16.0000 ± 0.5774 | -17.3520 to -14.6480 | 16.0000 |

**Table S3: Statistical analysis of male wing area variation for varying illuminance using Kruskal-Wallis test with Dunn's multiple comparisons test. Related to Figure 3C.**

| Comparison | Adjusted P-value | Mean rank difference |
|---|---|---|
| 5 vs 0 (Dark) | 0.039810 * | 15.2879 |
| 5 vs 3 | 0.04769 * | 15.8545 |
| 5 vs 10 | 0.001059 ** | 18.1818 |
| 5 vs 50 | 0.026910 * | 15.3117 |

**Table S4: Statistical analysis of male wing area variation for 5 lux/ 12 hrs vs 5 lux/ 1min illuminance using Unpaired t-test. Related to Figure 3D.**

| t | DF | p-value (two-tailed) | Difference Between Means (A - B) ± SEM | 95% Confidence Interval | Cohen's *d* |
|---|---|---|---|---|---|
| 5.4216 | 43.0000 | <0.0001 **** | -0.09919 ± 0.01829 | 0.06229 to 0.1361 | -1.8806 |

**Table S5: Statistical analysis of female wing area variation for varying illuminance using Kruskal-Wallis test with Dunn's multiple comparisons test. Related to Figure 3C.**

| Comparison | Adjusted P-value | | Mean rank difference |
|---|---|---|---|
| 5 vs 0 (Dark) | 0.0019810 | *** | 18.7921 |
| 5 vs 3 | 0.03769 | * | 15.4515 |
| 5 vs 10 | 0.000105 | **** | 18.9889 |
| 5 vs 50 | 0.0026910 | * | 15.1785 |

**Table S6: Statistical analysis of female wing area variation for 5 lux/ 12 hrs vs 5 lux/ 1min illuminance using Kruskal-Wallis test with Dunn's multiple comparisons test. Related to Figure 3D.**

| t | DF | p-value (two-tailed) | Difference Between Means (A - B) ± SEM | 95% Confidence Interval | Cohen's *d* |
|---|---|---|---|---|---|
| 3.7343 | 35.0000 | 0.000668 *** | -0.1290 ± 0.03456 | 0.05889 to 0.1992 | -1.4309 |

**Table S7: Statistical analysis of global (tissue wide) apoptosis for varying light illuminance levels using Kruskal-Wallis test with Dunn's multiple comparisons test. Related to Figure 4B.**

| Comparison | Adjusted P-value | Mean rank difference |
|---|---|---|
| Control: 0 lux vs 5 lux/ 1min | 0.2641 ns | -8.0000 |
| Control: 0 lux vs 5 lux/12 hrs | 0.001936 ** | -16.0000 |
| Control: 0 lux vs 100 lux/12 hrs | <0.0001 **** | -24.0000 |

**Table S8: Statistical analysis of compartment-specific (anterior vs posterior) apoptosis for varying light illuminance levels using Kruskal-Wallis test with Dunn's multiple comparisons test. Related to Figure 4C.**

| Comparison | Adjusted P-value | | Mean rank difference |
|---|---|---|---|
| **5 lux/1min (A) vs P, 5 lux/1min (P)** | **0.009102** | ** | **-24.0000** |
| 5 lux/1min (A) vs 5 lux/12 hrs (A) | 1.0000 | ns | -8.6250 |
| 5 lux/1min (A) vs 5 lux/12 hrs (P) | <0.0001 | *** | -32.0000 |
| 5 lux/1min (A) vs 100 lux/12 hrs (A) | 0.4209 | ns | -15.3750 |
| 5 lux/1min (A) vs 100 lux/12 hrs (P) | <0.0001 | **** | -40.0000 |
| P, 5 lux/1min (P) vs 5 lux/12 hrs (A) | 0.4209 | ns | 15.3750 |
| P, 5 lux/1min (P) vs 5 lux/12 hrs (P) | 1.0000 | ns | -8.0000 |
| P, 5 lux/1min (P) vs 100 lux/12 hrs (A) | 1.0000 | ns | 8.6250 |
| P, 5 lux/1min (P) vs 100 lux/12 hrs (P) | 0.3341 | ns | -16.0000 |
| **5 lux/12 hrs (A) vs 5 lux/12 hrs (P)** | **0.012510** | * | **-23.3750** |
| 5 lux/12 hrs (A) vs 100 lux/12 hrs (A) | 1.0000 | ns | -6.7500 |
| 5 lux/12 hrs (A) vs 100 lux/12 hrs (P) | 0.0001108 | *** | -31.3750 |
| 5 lux/12 hrs (P) vs 100 lux/12 hrs (A) | 0.2632 | ns | 16.6250 |
| 5 lux/12 hrs (P) vs 100 lux/12 hrs (P) | 1.0000 | ns | -8.0000 |
| **100 lux/12 hrs (A) vs 100 lux/12 hrs (P)** | **0.006526** | ** | **-24.6250** |

**Table S9: Statistical analysis of global (tissue-wide) mitosis for varying light illuminance levels using Kruskal-Wallis test with Dunn's multiple comparisons test. Related to Figure 5B.**

| Comparison | Adjusted P-value | | Mean rank difference |
|---|---|---|---|
| Control: 0 lux vs 5 lux/1 min | <0.0001 | **** | -24.0000 |
| Control: 0 lux vs 5 lux/12 hrs | 0.001059 | *** | -18.5833 |
| Control: 0 lux vs 100 lux/12 hrs | 0.003281 | *** | -16.4167 |

**Table S10: Statistical analysis of compartment-specific (anterior vs posterior) mitosis for varying light illuminance levels using Kruskal-Wallis test with Dunn's multiple comparisons test. Related to Figure 5C.**

| Comparison | Adjusted P-value | | Mean rank difference |
|---|---|---|---|
| **5 lux/1 min (A) vs 5 lux/1 min (P)** | **1.0000** | **ns** | **-3.6667** |
| 5 lux/1 min (A) vs 5 lux/12 hrs (A) | 0.0006234 | *** | 24.9167 |
| 5 lux/1 min (A) vs 5 lux/12 hrs (P) | 1.0000 | ns | 8.1667 |
| 5 lux/1 min (A) vs 100 lux/12 hrs (A) | 0.05490 | ns | 17.6667 |
| 5 lux/1 min (A) vs 100 lux/12 hrs (P) | 0.3310 | ns | 13.9167 |
| 5 lux/1 min (P) vs 5 lux/12 hrs (A) | <0.0001 | *** | 28.5833 |
| 5 lux/1 min (P) vs 5 lux/12 hrs (P) | 0.7739 | ns | 11.8333 |
| 5 lux/1 min (P) vs 100 lux/12 hrs (A) | 0.006742 | ** | 21.3333 |
| 5 lux/1 min (P) vs 100 lux/12 hrs (P) | 0.05735 | ns | 17.5833 |
| **5 lux/12 hrs (A) vs 5 lux/12 hrs (P)** | **0.08796** | **ns** | **-16.7500** |
| 5 lux/12 hrs (A) vs 100 lux/12 hrs (A) | 1.0000 | ns | -7.2500 |
| 5 lux/12 hrs (A) vs 100 lux/12 hrs (P) | 1.0000 | ns | -11.0000 |
| 5 lux/12 hrs (P) vs 100 lux/12 hrs (A) | 1.0000 | ns | 9.5000 |
| 5 lux/12 hrs (P) vs 100 lux/12 hrs (P) | 1.0000 | ns | 5.7500 |
| **100 lux/12 hrs (A) vs 100 lux/12 hrs (P)** | **1.0000** | **ns** | **-3.7500** |

**Table S11: Statistical analysis of area tumor over area of whole wing disc using Kruskal-Wallis test with Dunn's multiple comparisons test. Related to Figure 6F.**

| Comparison | Adjusted P-value | Mean rank difference |
|---|---|---|
| No RasV12 vs 0 lux | 0.004696 ** | -12.2500 |
| No RasV12 vs 5 lux/1 min | >0.9999 ns | -2.9167 |
| No RasV12 vs 5 lux/12 hrs | 0.1878 ns | -7.5000 |
| No RasV12 vs 100 lux/12 hrs | 0.02318 * | -11.2500 |
